# Loss of TTC5 impairs cortical development, disrupts tubulin homeostasis and progenitor mitosis in vivo

**DOI:** 10.64898/2026.11.25.768198

**Authors:** Zhangshen Li, Li Ma, Hasib A. Riaz, Qing Wu, Michael Schmidt, Morgan Fleishman, Eric M. Morrow

## Abstract

**Objective:** Loss-of-function mutations in the multifunctional scaffolding protein tetratricopeptide repeat domain 5 (TTC5) cause Neurodevelopmental Disorder with Cerebral Atrophy and variable Facial Dysmorphism (NEDCAFD), a recently described neurogenetic syndrome. TTC5 autoregulates tubulin translation in vitro; however, the role of TTC5 in brain development in vivo is poorly understood. Here, we define the function of TTC5 in neocortical development.

**Methods:** We established a mouse model carrying a recurrent patient nonsense mutation (*TTC5*^R263*/R263*^) causing NEDCAFD, and investigated cortical progenitor development, tubulin homeostasis, microtubule function, and neuronal morphogenesis using in vivo, in vitro, and transcriptomic approaches.

**Results:** The *TTC5^R263*^* mutation was a strong loss-of-function allele, with homozygous newborns exhibiting neonatal lethality, reduced brain size, and loss of cortical neurons. These developmental abnormalities arise from an early and persistent loss of neural progenitor expansion and impaired neuronal differentiation during corticogenesis, associated with defective mitotic progression, altered cell fate determination, and increased DNA damage. We further investigate TTC5 as a regulator of tubulin homeostasis during cortical development in vivo. Loss of TTC5 reduced tubulin mRNA and protein levels. Furthermore, TTC5^R263*/R263*^ primary neurons exhibit reduced microtubule growth dynamics and defective neuronal arborization. Transcriptomic analyses of TTC5^R263*/R263*^ developing cortex identify differentially expressed genes and pathways involved in neurodevelopmental processes, including coordinated dysregulation of cell-cycle regulators and a multitude of microtubule-associated gene pathways.

**Interpretation:** TTC5 is essential for cortical progenitor expansion, mitotic progression, and tubulin homeostasis during cortical development. These findings establish a mechanistic framework for NEDCAFD pathogenesis in vivo.

## Introduction

Intellectual disability (ID) is highly heterogeneous with a global prevalence of 1-3%.^1^ Autosomal recessive ID (ARID) accounts for about 10% of all ID and 20-25% of monogenic ID in families without recent shared ancestry.^2,3^ The risk of ARID is significantly higher in consanguineous families, with autosomal recessive genetic mutations being the leading genetic cause of ID in populations with high parental consanguinity.^2,4^ Large ARID studies over the past decade have identified approximately 400 candidate genes.^2,4–8^ These studies have contributed to the genetic diagnosis of ID for families across the world. In addition, advances in the genetics of ARID have provided important insight into the genetic architecture of human intellectual function, as well as mechanisms of human brain development.

In recent studies of ARID, putative loss-of-function mutations were identified in the gene encoding the multifunctional scaffolding protein tetratricopeptide repeat domain 5 (TTC5).^4,9–11^ In one study, *TTC5* was one of four genes with multiple identified allelic mutations that have not been previously associated with any disease.^4^ The new TTC5-associated gene syndrome has been termed the Neurodevelopmental Disorder with Cerebral Atrophy and variable Facial Dysmorphism (NEDCAFD). In 19 published patients, the NEDCAFD syndrome presents with intellectual disability (100%), severe speech impairment (100%), motor delay (100%), seizures (26%), microcephaly (63%) and facial dysmorphism (88%). On MRI, structural brain differences include cerebral atrophy (75%), enlarged lateral ventricles (75%), corpus callosum abnormalities (69%), and simplified gyral patterning (75%).^9–11^ To date, there are no studies investigating the mechanisms underlying neurodevelopmental abnormalities in NEDCAFD due to apparent loss- of-function mutations in TTC5.

TTC5 (also known as STRAP, Stress-Responsive Activator of p300) is a 440-amino acid protein consisting of a tandem series of six tetratricopeptide repeat (TPR) motifs, which are structural modules mediating protein-protein interactions.^12,13^ A TPR domain comprises multiple repeats of 34 amino acid units forming a series of repeating antiparallel α helices due to their parallel packing.^14^ This overall super-helix structure forms binding pockets for diverse ligands with different surface residues for specific ligand interaction, and thereby, TPR motif-containing proteins are involved in diverse biological processes including cell cycle control, stress response, and chaperone functions.^14–19^ In addition to the TPR domains, TTC5 contains an oligonucleotide binding (OB) fold at the C-terminal, which is a functional module widely implicated in genomic maintenance, especially in DNA damage response.^20^ Indeed, TTC5 augments p53 response under cellular stress by downregulating MDM2 activity. Stress-responsive accumulation of TTC5 promotes the assembly of a p53 coactivator complex, through direct interaction with cofactors JMY (via TPR motifs 1–3 or 3–5) and p300 (via TPR motifs 5, 6, and the OB fold).^13,21,22^ TTC5 is regulated by DNA damage signalling pathways, where ATM (ataxia telangiectasia mutated) phosphorylation prompts nuclear accumulation of TTC5 by impeding nuclear export and CHEK2 (Checkpoint kinase 2) phosphorylation augments nuclear TTC5 stability.^16^ Additionally, TTC5 modulates the activity of nuclear hormone receptors, including the glucocorticoid receptor (GR). TTC5 may stabilize GR by preventing its ubiquitination and subsequent proteasomal degradation, thereby influencing GR-mediated transcriptional responses. This regulation is particularly significant under stress conditions, where TTC5 can alter GR activity in a gene-specific manner.^17^

Importantly, TTC5 has been proposed to play a critical role in the autoregulation of tubulin synthesis.^10,23–25^ TTC5 binds to ribosomes translating tubulin mRNAs and recognizes the nascent tubulin polypeptides. Upon detecting excess soluble tubulin, TTC5 recruits the protein SCAPER (S-phase cyclin A associated protein residing in the ER), which in turn engages the CCR4-NOT deadenylase complex via its CNOT11 subunit, leading to the degradation of tubulin mRNAs.^23^ This mechanism ensures the maintenance of tubulin homeostasis, crucial for microtubule function.^23–27^ Proper tubulin function is essential for brain development, and multiple genetic abnormalities of tubulin have been implicated in developmental brain disorders, namely tubulinopathies, which are neurological disorders characterized by malformations of cortical development (MCD) resulting from defective cortical neuronal proliferation, migration, and postmigrational development.^28–30^ Microtubules are assembled by αβ-tubulin heterodimers, and mutations in genes encoding different tubulin isotypes lead to impaired microtubule polymerization and altered microtubule dynamics. Such perturbations result in disruptions in major cellular processes in neuronal development, including mitosis and cytokinesis of progenitors, differentiation and positioning of new-born neurons, morphogenesis and pathfinding of axons and dendrites, and intracellular cargo trafficking.^30–34^ A possible connection between NEDCAFD and tubulinopathies has been proposed, with clinical features resembling tubulinopathy-associated brain malformations.^10^

The diverse functions of TTC5 highlight its critical role in cellular homeostasis and potentially in development; however, to date there have been no studies investigating the function of TTC5 in the developing nervous system in vivo. Given the critical role of TTC5 in the developing brain, as exemplified by mutations in novel syndrome NEDCAFD, we mutated the TTC5 protein in mouse, including a mutation found in patients with NEDCAFD, i.e. the nonsense variant p.(Arg263Ter). In our studies, we define this patient-derived mutation as a loss-of-function mutation and subsequently discover that TTC5 is required for mouse neonatal survival. *TTC5^R2^*^63*\*/R263\**^ mutation leads to mitotic progenitor defects in embryonic cortical neural precursors causing reduced brain size and abnormal cortical development. Transcriptome analysis in neonatal *TTC5^R263*/R263*^* neocortex demonstrates a dysregulation of tubulin mRNA abundance. Further, *TTC5^R263*/R263*^* mutation affects microtubule protein abundance and dynamics in cortical neurons associated with neuronal morphogenesis defects. In summary, our studies demonstrate that TTC5 is essential for progenitor mitosis and proper development of the cerebral cortex and highlight the role of TTC5 in sustaining tubulin homeostasis in the developing brain in vivo, providing insights into the pathogenesis of NEDCAFD.

## Materials and Methods

### Mice

All animal work was conducted under the guidelines of the Center for Animal Resources and Education (CARE) with a protocol (IACUC 21-09-0012), approved by the Brown University Institutional Animal Care and Use Committee (IACUC). All experimental procedures were consistent with the US National Institutes of Health Guide for the Care and Use of Laboratory Animals (National Research Council, 8th edition). Animals were maintained on a 12/12-hour light/dark cycle with ad libitum access to food and water.

### Mouse model generation

The Transgenic and Genome Editing Facility at Brown University generated the TTC5*^R263*/R263*^* mouse model using CRISPR/Cas9-mediated genome editing on C57BL/6J background, as described herein. A nonsense point mutation was introduced in mouse Ttc5 exon 7 of 10 (transcript ENSMUST00000006451.9), matching the consensus human pathogenic variant *c*.787C>T; p.(Arg263Ter).

Recombinant Cas 9 protein, single guide RNA with the spacer sequence (GCAA***CGA***GAACAGCAACTCC), and a single-stranded oligodeoxynucleotide (ssODN) as a homology-directed repair (HDR) template (GAGGAGAGTTATGGGGAGGCCCTTGAGGGTTTCTCTCAGGCTGCAGCGCTGGACCCTGT GTGGCCAGAGCCCCAGCAA***tGA***GAACAGCAACTCCTcGAATTCCTCAGTAGGCTAACCAGC CTCCTGGAGAGCAAGGTAAAGACGCGTAAGGGCTGAAGCTGTCAGTCAG) were mixed and microinjected into mouse embryos to generate founder mice. Two independent mosaic F0 founders with the targeted R263* mutation were generated and confirmed by Sanger sequencing (Transgenic and Genome Editing Facility at Brown University). Founders were backcrossed to wild-type mice on C57BL/6J background for three generations prior to phenotyping, and animals derived from both founders were included in the phenotypic analyses. The presence of the mutation was confirmed by PCR amplification (Forward, 5′-TCAGACTTTGTGAGGTAGCTTCT-3′; Reverse, 5′-AAGGGAAAGTGAGACTTACAAGGG-3′) followed by Sanger sequencing, or by Tetra-primer Amplification Refractory Mutation System (ARMS) PCR genotyping (Forward inner primer (T allele): GTGTGGCCAGAGCCCCAGCCAT; Reverse inner primer (C allele): GGAATTCCAGGAGTTGCTGTTCGCG; Forward outer primer (5’ - 3’): GAAGAGCGAGTTCCGTTTGACCCCA; Reverse outer primer (5’ - 3’): ACTGATAGCGCCCATCACCACAGGG). Tetra-primer ARMS PCR was performed as described in Medrano et al., 2014 with modifications.^35^ Detailed primer information can be seen in **Supplementary Table 9**.

### Tissue preparation, immunohistochemistry and antibodies

Embryonic and postnatal day 0 (P0) whole heads were fixed in 4% paraformaldehyde by immersion for 24 hours and 96 hours at 4°C, respectively. Whole head tissues were then washed with PBS and cryopreserved in sucrose solutions in PBS from 10% to 30% at 4°C until tissue sank. Whole heads were then embedded in OCT freezing media and stored at -80°C. Coronal cryosections of 20*μ*m (for embryonic tissue) or 30μm (for P0 brain tissue) were generated using a Leica Cryostat and mounted sections were stored at -80°C. Heat-induced antigen retrieval (HIER) was carried out by immersing sections in sodium citrate buffer (10mM Sodium citrate, 0.05% Tween 20, pH 6.0) at sub-boiling temperature for 10 min and cooled down to room temperature.

To detect cell proliferation, 5-ethynyl-2^’^-deoxyuridine (EdU) was injected intraperitoneally into pregnant females at E13.5 (10 *μ*g/g of body weight). Embryos were harvested 24 hours after injection and brain tissues were processed as described above. EdU detection assay was performed using the Click-iT™ Plus EdU Cell Proliferation Kit for imaging (Thermo Fisher, C10638).

Immunohistochemistry was conducted by permeabilizing sections in 0.5% Triton X-100 in PBS for 30 min, blocking in 10% normal goat serum and 0.25% Triton X-100 in PBS for 2 hours, followed by incubation with primary antibodies in 2% goat serum and 0.25% Triton X-100 in PBS at 4°C overnight. Sections were then washed in PBS and incubated with corresponding fluorescence-conjugated secondary antibodies for 2 hours at room temperature in the same solution. Coverslips were mounted on slides using Fluoromount-G (Southern Biotech, 0100-01) following washes in PBS.

The primary antibodies used for immunohistochemistry at specific dilutions were as follows: mouse anti-Satb2 (1:500) (Abcam, ab51502); rat anti-Ctip2 (1:500) (Abcam, ab18564); rabbit anti-Tbr1 (1:500) (Abcam, ab31940); mouse-anti Ki67 (1:400) (BD Biosciences, 550609); rat anti-nestin (1:200) (DSHB, Rat-401); rabbit anti-Dcx (1:400) (Abcam, ab18723); rabbit anti-Pax6 (1:400) (Abcam, ab195045); rabbit anti-Tbr2 (1:200) (Abcam, ab23345); rat anti-phospho-histone H3 (1:800) (Abcam, ab10543); mouse anti-aurora kinase B (AURKB) (1:400) (BD Biosciences, 611083); rabbit anti-cleaved Caspase-3 (1:200) (Cell Signaling, 9661); mouse anti-γ-H2AX (1:200) (Cell Signaling, 80312). Secondary antibodies used include Alexa Fluor 488-, 555-, or 647-conjugated to goat anti-mouse, rabbit, or rat IgG at 1:500 (Invitrogen). Nuclei were stained and visualized with Hoechst33342 (1:1500) (Invitrogen).

### TUNEL assay

TUNEL assay was performed using the Click-iT™ Plus TUNEL Assay Kits for In Situ Apoptosis Detection (Thermo Fisher, C10617). Tissue sections were processed as described in the section above and TUNEL labeling was performed following the manufacturer’s instructions. Positive controls were generated by treating brain sections with 1 unit of DNase I for 30 min at room temperature. Nuclei were visualized with Hoechst33342 (1:1500) (Invitrogen).

### Image acquisition and data analysis

All fluorescence images were acquired using an Olympus FV3000 confocal microscope or a Nikon Spatial Array Confocal (NSPARC) microscope with 20x or 60x objective lens. For each experiment, at least three coronal serial brain sections from at least three different animals per genotype were analyzed. Quantitative measurements of cell numbers were performed on images of comparable sections from each genotype using ImageJ processing program (NIH).

For cortical neuronal analysis at P0, regions of defined thickness were selected in each layer (layers II-IV, 100*μ*m; layer 5, 100*μ*m; layer VI, 150*μ*m) of the cortical plate from images of equal width (60x, 200*μ*m) and cortical cell numbers were quantified on at least 3 consecutive z-positions from a 30μm z-stack. The average percentage of marker-positive neurons were calculated relative to the total number of nuclei in each region by ImageJ in consecutive z-positions. Neural stem cells at P0 were quantified as the number of nestin and Ki67 double-positive cells in a 200*μ*m x 200*μ*m region in the ventricular zone of mid-anterior cortex.

For mitotic spindle orientation analysis, angles of cleavage planes in dividing neuronal progenitors in the ventricular zone were measured relative to the ventricular surface, on serial z-positions from a 20*μ*m z-stack, from anaphase and telophase cells with clearly defined mitotic spindles and two daughter cells.

For γ-H2AX foci analysis, the number of foci per nucleus was measured in a 200*μ*m x 200*μ*m region in the ventricular zone from a 20*μ*m z-stack. Regions of interest (ROIs) were defined around individual nuclei visualized by Hoechst staining and the number of nuclear foci were counted after universal thresholding.

### Bulk RNA Sequencing and analysis

P0 TTC5*^+/+^* and *TTC5^R263*/R263*^* cortical tissues (n = 5) were dissected (hippocampus removed), and total RNA was isolated from the right cortices using RNeasy Mini Kit (Qiagen 74104) according to the standard protocol. RNA extraction quality was verified by the rRNA Ratio [28 s/ 18 s] and RNA integrity number (RIN) and sequencing of poly-(A) enriched RNA was carried out by Azenta Life Sciences.

A FastQC report was generated to confirm read quality. The STAR-StringTie-DESeq2 pipeline was conducted with an average coverage of 46 million reads per sample. A threshold of padj =0.05 was used for defining differentially expressed genes (DEGs).

### snRNAseq library preparation

Cortices from WT (n=4) and MUT (n=4) mice were dissected from postnatal day 0 pups. The WT group comprised one female and three males, and the MUT group comprised two females and two males. Nuclei were isolated following the manufacturer’s instruction from Nuclei Isolation Kit (Millipore sigma NUC201). Briefly, the cortex was minced in a Dounce homogenizer and filtered through a 40 *μ*m cell strainer. Nuclei suspensions were centrifuged at 30,000g for 45 min at 4°C for the sucrose purification step.^36^ The nuclei pellets were resuspended in nuclei storage buffer, and nuclei were counted using a hemocytometer. Single nuclei were captured using the 10x Genomics Chromium Controller with the Chromium Next GEM Single Cell 3’ Reagent Kits v4.1, and cDNA and libraries were subsequently prepared following the manufacturer instructions (10x Genomics). All samples were processed and sequenced together. Because of the small number of animals per sex, sex was not included in the analysis, and sex-linked genes (Xist, Tsix, Ddx3y, Eif2s3y, Uty, and Kdm5d) were removed from differential expression analysis.

### snRNAseq data analysis

Sequence alignment was performed against the Mus musculus reference genome (GRCm39-2024) using Cell Ranger software (9.0.1) provided by 10X genomics. CellBender was used to remove ambient RNA, and the processed data were subsequently analyzed using Seurat (v5).^37^ To remove low-quality cells, nuclei expressing fewer than 200 genes or those with more than 6000 genes or with mitochondrial content greater than 10% were excluded. We performed Harmony Integration through Seurat by using the Integrate Layers function, followed by the standard Seurat clustering pipeline for clustering and downstream analysis.^38^ Cell type identities were determined by the expression of canonical marker genes. We identified Layer 1 neurons by Reln and Trp73 expression, Layer 2-3 neurons by Cux1 and Satb2, Layer 4 neurons with Rorb and Unc5d, Layer 5-6 neurons with Tbr1 and Foxp2 and immature excitatory neurons with Dcx, Unc5d and Sema3c. Inhibitory neurons were identified with Gad1 and Gad2, Neuronal progenitors with Eomes, Proliferating and glial cells with Mki67, Microglia with Cx3cr1 and Csf1r, Astrocytes with Aldh1l1, OPC and Oligodendrocytes by Pdgfra and Sox10, Endothelial cells with Pecam1 and Flt1, Pericytes with Rgs5 and Pdgfrb and VLMCs (vascular and leptomeningeal cells) with Dcn.^39–41^ The cell type initially identified as “Proliferating and glia” was further re-clustered, and cell cycle phase scores (S and G2/M) were calculated using Seurat’s CellCycleScoring function. Subclusters were further identified as three subtypes: Dlx+ inhibitory-lineage progenitors (Dlx1, Dlx2, Dlx5, Dlx6os1, Sp9), radial glia/glial progenitors (Fabp7, Sparcl1, Tnc, Aldh1l1, Slc1a3), and postmitotic neurons (Syt1, Camk2b, Scn2a) using Seurat’s AddModuleScore function.^37^

Differential expression between genotypes was assessed separately for each cell type using a pseudobulk approach, with individual animals treated as the unit of replication. For each animal, raw counts were aggregated across all nuclei within a given cell type.^42,43^ Animal level cell type profiles containing fewer than 20 nuclei were excluded, and a cell type was included in the analysis only if at least three animals per genotype remained after filtering. In addition to the sex-linked genes Xist, Tsix, Ddx3y, Eif2s3y, Uty, and Kdm5d, genes that did not have at least 10 counts in at least three animals were removed. Differential expression between genotypes was tested with DESeq2 using the Wald test.^44^ Sex was not included as a covariate because the number of animals per sex and genotype was too small to estimate it reliably.

To complement the pseudobulk analysis with a nucleus level analysis, we used the negative binomial gamma mixed model (NEBULA) on raw counts within each cell type.^45^ Genotype was modeled as a fixed effect and animal as a random effect, and total counts per nucleus were used to correct for differences in sequencing depth. Genes detected in fewer than 10% of nuclei, sex-linked genes, and genes for which the model did not converge were excluded. P values were adjusted within each cell type using the Benjamini-Hochberg method, and genes with an adjusted p value < 0.05 were considered significant. Gene set enrichment analysis (GSEA) was performed per cell type with fgsea using hallmark gene set. The pathways with padj < 0.1 were called significant.

### Tissue preparation and protein assay by western blotting

P0 mouse brain tissues were homogenized and lysed in Radioimmunoprecipitation assay buffer containing 50mM Tris-HCl, pH 7.4, 150mM NaCl, 1mM EDTA, 1% Triton X-100, 1% sodium deoxycholate and 0.1% SDS supplemented with protease inhibitor cocktail (Roche, 05892970001) and phosphatase inhibitor (Roche, 04906837001) for 1hr on ice and were centrifuged at 13,200 rpm for 15 min at 4°C. The supernatant protein concentration was measured by bicinchoninic acid (BCA) assay using the Pierce BCA Kit (Thermo fisher Scientific, 23225).

Tissue lysates were then boiled in sample buffer at 95 °C for 5 min before loading onto 4-12% SDS-PAGE gels (Novex #NP0321Box). Following separation of proteins by electrophoresis, gels were then transferred to nitrocellulose membranes (Novex #LC2000) and blotted with anti-TTC5 antibody (Proteintech, 26112-1-AP), Tubulin antibodies (mouse-anti-β3-Tubulin (Cell Signaling, 4466S), Rabbit-anti-β3-Tubulin (Cell Signaling, 5568S), Mouse-anti-acetyl-α-Tubulin (Sigma, T7451), Rabbit-anti-α-Tubulin (Sigma, T6074), Mouse-anti-γ-Tubulin (Sigma, T6557), pan β Tubulin (C-10) (Santa Cruz, sc-398937)) overnight at 4°C. Western blots were performed using standard procedures^46,47^ and were analyzed with the Li-CoR Odyssey Imaging System. Tubulin bands were quantified by densitometry using Li-CoR software and normalized to GAPDH.

### Primary neuronal culture and transfection

Primary neurons from P0 mouse cortices were dissociated with papain as described before.^48^ All plates/dishes were pre-coated with 1 mg/ml poly-D-lysine at 37 °C for at least 2hr ahead of cell seeding.

For microtubule dynamics and behavior live imaging, cortical neurons cultured on 35 mm dish were transfected with EB3-tdTomato construct (addgene Plasmid #50708) using Lipofectamine 2000 (Invitrogen) according to manufacturer’s instructions.

### Continuous live-cell imaging and neurite outgrowth analysis

IncuCyte S3 Live-Cell Analysis System (Essen BioScience) was used for continuous live-cell imaging. 6 replicate wells of isolated neuronal cells per pup were seeded into 96-well plates. Plates were then placed into the IncuCyte S3 Live-Cell Analysis System at 1 DIV, followed by automated continuous live-cell imaging at intervals of 6 h with a 10× objective lens for 14∼15 DIV in total. Phase-contrast images were collected. The recorded images were then analyzed using the automated NeuroTrack image acquisition module of the IncuCyte Zoom software. The NeuroTrack algorithm automatically defines cell bodies and neurite branches to calculate and quantify neurite length and neurite branch points per cell body cluster. Neurite length and neurite branch points data were normalized to their respective values at time 0 h and are presented as fold changes relative to baseline (time 0 h = 1).

### Microtubule polymerization and co-sedimentation assay

TTC5 microtubule polymerization and co-sedimentation assay was performed as described previously^49^ with some modifications. In brief, P0 mouse brain tissues were dissected and homogenized in ice-cold BRB80 buffer (80 mM PIPES / KOH, 1mM EGTA, 1mM MgCl2, pH 6.8) supplemented with protease inhibitor cocktail (Roche, 05892970001) and phosphatase inhibitor (Roche, 04906837001). Crude tissue lysates were clarified by centrifugation at 14,000 rpm for 15 mins at 4°C, and then clarified lysates were centrifuged at 18,000g for 1hr at 4°C. Supernatants (S1 fraction) at equal protein concentrations (about 400*μ*g) were added at final concentration with 1 mM GTP (Sigma) and 20% glycerol. These mixtures were incubated for 1hr at 37°C to allow tubulin to polymerize into microtubules, and then were centrifuge at 20,000 x g for 1hr at 25°C. This yielded a pellet containing microtubules and associated proteins (P2 fraction) and a soluble supernatant (S2 fraction). The P2 fraction was then re-suspended and depolymerized completely in BRB80 buffer. All fractions were denatured with loading buffer and analyzed by SDS-PAGE and western blot.

### MT Dynamics Assays in Cultured Cortical Neurons

Neurons dissociated from mouse P0 cortices were grown on 35mm dishes (D35-14-1.5N, Cellvis). On DIV 2, the cells were transfected with EB3-TdTomato construct for 24hr and live-cell imaging of EB3 comets was performed in culturing medium using Nikon Spatial Array Confocal (NSPARC) super resolution microscope with a temperature-controlled (37 °C) CO_2_ incubator (6 sec/frame for 3 min). Comet movements were analyzed by ImageJ Plugin MTrackJ for motion tracking and analysis. Parameters describing MT dynamics were defined as follows: comet growth rate is the growth length/comet lifetime (*μ*m/sec), comet lifetime is the duration of growth in seconds, and comet growth length is the comet movement length in micrometers.^50,51^ Image acquisition and quantification were blinded to the genotype.

### Statistical analysis

Statistical analyses were performed with unpaired two-tailed t-test with Welch’s correction, one-way ANOVA with Tukey’s/Dunnett’s multiple comparisons test, or Mann–Whitney test as indicated in figure legends. Graphs were plotted using GraphPad Prism software. Data are presented as mean ± SD with significance levels * for *P*<0.05, ** for *P*<0.01, *** for *P*<0.001 and **** for *P*<0.0001.

## Results

### Patient *TTC5^R263*/R263*^* mutation causes loss of function and demonstrates that TTC5 is required for neonatal survival in mouse

For genetic construct validity for the NEDCAFD syndrome, we initially generated a mutation in mouse that is homologous to mutations found in NEDCAFD patients. The human *c*.787C>T; p.(Arg263Ter) (NM_138376.3) homozygous mutation was first reported by Rasheed et al. (2021).^9^ Patients with this mutation exhibited phenotypes including severe intellectual disability, non-verbal status, hypotonia, facial dysmorphism, and abnormal brain MRI with cerebral atrophy, enlarged lateral ventricles, corpus callosum atrophy, simplified gyral pattern.^9^ This mutation is located in the linker domain which is between TTC5 TPR6 and OB domain. To understand the mechanism by which this mutation causes NEDCAFD, we modelled this mutation in vivo in mice. We generated the TTC5 transgenic mouse line through pronuclear microinjection of CRISPR-Cas9 and gRNA, resulting in a nonsense mutation p.(Arg263Ter) on exon 7 of the *TTC5* allele on mouse chromosome 14 (**Fig. 1a**). The founder lines were generated in, backcrossed to and propagated in the C57/BL6J genetic background. In Tetra PCR, a band of 310bp generated by the forward inner T allele primer and reverse outer primer indicates a homozygous mutant genotype. A band of 420bp from the forward outer primer and reverse inner C allele primer indicates a wild-type genotype. The presence of both bands indicates a heterozygous genotype (**Fig. 1b**). Sanger sequencing of genomic DNA from tail biopsies confirmed the C>T mutation in the *TTC5* allele (**Fig. 1c**).

**Fig. 1.**
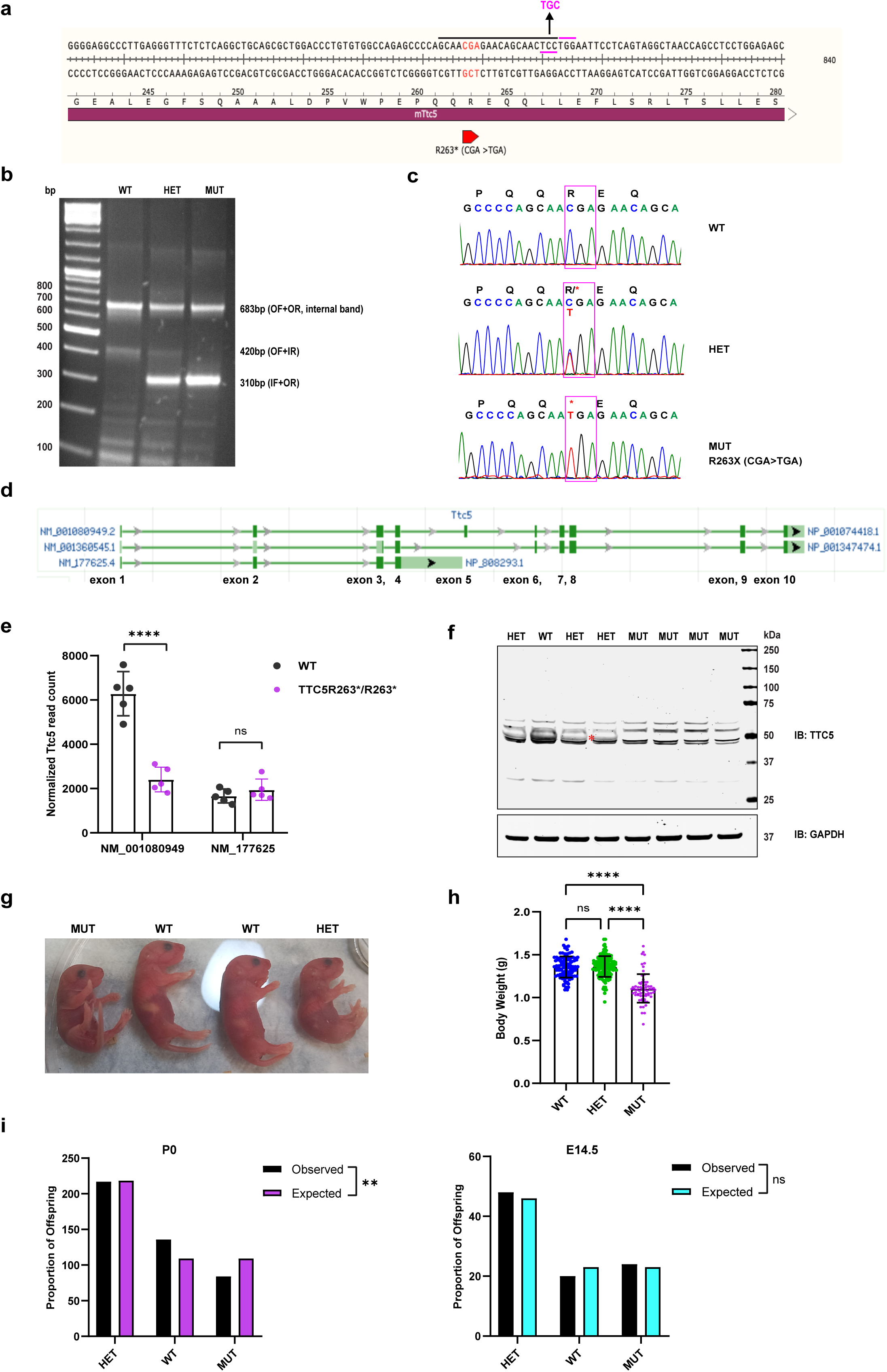
Generation and validation of patient-specific *TTC5^R263*/R263*^* mice. **a** Schematic of the CRISPR/Cas9 knock-in methodology used to generate the *c*. 787 C>T; R263* (NM_138376.3, NP_612385.2). Underlined depicts Guide RNA. Orange color indicates the protospacer adjacent motif (PAM) sequences. Red font indicates the mutated nucleotides. Magenta depicts the silent mutation region to prevent further cleavage from Cas9. **b** Tetra-primer ARMS-PCR genotyping was performed on tail lysates from *TTC5^+/+^*, *TTC5^+/R263*^*, and *TTC5^R263*/R263*^* mice. All conditions yield an internal band at the size of 683bp. 420bp band represents the wild type genotyping, and 310bp band depicts the mutant. The HET pups have both bands. **c** Sanger sequencing confirmed the predicted nonsense mutation CGA>TGA in homozygous *TTC5^R263*/R263*^*mice. **d** The diagram shows that *Ttc5* has three transcripts (NCBI reference: NM_001080949.2, NM_1360545.1 and NM_177625.4) in mouse. **e** Normalized read counts of *Ttc5* transcripts: NM_001080949.2, and NM_177625.4(NCBI) analyzed from Bulk RNA sequencing wild type and *TTC5^R263*/R263*^* mice. Two-way Anova with Bonferroni’s multiple comparisons test was used (****, *p* < 0.0001 for WT vs. MUT in NM_001080949; ns, *p* = 0.9862 for WT vs. MUT in NM_177625). **f** Western blotting was performed to determine TTC5 protein expression in *TTC5^+/+^*, *TTC5^+/R263*^*, and *TTC5^R263*/R263*^* mouse brains. GAPDH was used as a loading control. Red asterisk indicates the TTC5 band. Ladders were labelled on the right side of gel. **g** Representative pups’ pictures at postnatal day 0. From left to right, MUT, WT, WT and HET. **h** Graph shows the body weight of P0 pups from *TTC5^+/+^*(blue), *TTC5^+/R263*^* (green), and *TTC5^R263*/R263*^* (magenta) mice. One-way Anova with Tukey’s multiple comparisons test was used (ns, *p* = 0.9246 for WT vs. HET; ****, *p* < 0.0001 for WT vs. MUT and HET vs. MUT). Animal numbers: n=101 for WT, n=144 for HET and n=65 for MUT. **i** Mendelian inherence ratio analysis using Chi-square test for P0 pups (left graph) and E14.5 embryos (right graph). For P0 pups, observed animal HET=217, WT=136, MUT=84; for E14.5 embryos, observed animal HET=48, WT=20, MUT=24. (**, *p =* 0.002 for P0 pups, ns, *p =*0.5134 for E14.5 embryos comparison). The total number of observed pups is shown in black, and the expected numbers are shown in magenta (P0) and cyan (E14.5), respectively.

The mouse *Ttc5* gene has three mRNA transcripts, namely NM_001080949.2, NM_001360545.1 and NM_177625.4, which encode the protein isoforms NP_001074418.1, NP_001347474.1 and NP_808293.1, respectively (**Fig. 1d**). Clustal protein alignment showed that NP_001074418.1 encodes the full-length 440-amino acid TTC5 protein. NP_001347474.1 encodes 292 amino acids, which contain the patient nonsense mutation site but lacks the first 145 N-terminal amino acids. NP_808293.1 encodes only the N terminus (1-185 amino acids) of TTC5 protein and does not include the nonsense mutation site in exon 7 (**Supplementary Fig. 1**). Massively parallel, short-read RNA sequencing on mRNA from postnatal day 0 (P0) mouse brain cortices revealed a drastic reduction in *TTC5* mRNA level in *TTC5^R263*/R263*^* (mutant 2407 ± 558.0 normalized counts vs. control 6282 ± 998.5) of the transcript (NM_001080949.2) compared to the wild-type controls. The shortest 5’ transcript (NM_177625.4), that does not contain the nonsense mutation, was expressed at comparable levels between the wild type and *TTC5^R263*/R263*^* although at a low level (mutant 1946 ± 481.0 vs. control 1662 ± 311.4) (**Fig. 1e**). We validated a TTC5 antibody raised to the N-terminus of the protein. Western blotting protein assay using this antibody to the N-terminus of TTC5 showed that TTC5 was detected with the highest level in wild type, to a lesser extent, in heterozygous mice, and there was loss of TTC5 protein expression in the *TTC5^R263*/R263*^* mutant (**Fig. 1f**). We did not detect truncated protein in the homozygote or heterozygote suggesting that a truncated protein is not generated, reflecting the likelihood that the *TTC5^R263*/R263*^* mutation leads to nonsense-mediated mRNA decay in the mouse brain.

In patients with TTC5 mutations, postnatal growth retardation has been reported.^4,10^ Interestingly, newborn (P0) *TTC5^R263*/R263*^* mice consistently displayed lower body weight (mutant 1.11 ± 0.166g vs. control 1.36 ± 0.123g) (**Fig. 1g, h**) indicating a prenatal developmental defect associated with this mutation. In addition, by monitoring *TTC5^R263*/R263*^*mutant pups, we noticed that some were either dead or appeared unhealthy at birth. Among 84 homozygous *TTC5^R263*/R263*^* pups from 22 breeding pairs, 26 were born dead, and 8 unhealthy, while the remaining 50 pups survived birth; however, no *TTC5^R263*/R263*^* pups survived past P1 (**Supplementary Fig. 2a-b**). These findings suggest that the *TTC5^R263*/R263*^* mutation-related lethality may begin during embryonic development and also involves newborn lethality. The Mendelian inheritance ratio analysis was then performed using a Chi-square test at two timepoints, P0 and embryonic day 14.5 (E14.5). For P0 pups, we collected a total of 437 pups from 66 litters; and for E14.5 embryos, we collected 92 embryos from 13 females. As shown in **Fig. 1i**, for P0, while the number of heterozygous individuals matched the expected value (observed 49.66% vs. expected 50%), there were fewer homozygous mutants (observed 19.22% vs. expected 25%) and more wild-type individuals (observed 31.12% vs. expected 25%) than predicted (*p =* 0.002 for P0 pups comparison). In contrast, the E14.5 embryos showed no significant difference between the observed and expected genotype distributions across all three genotypes (ns, *p =*0.5134 for E14.5 embryos comparison). These results suggest that Mendelian inheritance patterns were maintained in E14.5 embryos but disrupted at P0. This implies that lethality associated with the *TTC5^R263*/R263*^* mutation occurs after embryonic stage E14.5, during late embryonic development and the early postnatal period. In summary, we generated and validated *TTC5^R263*/R263*^* mice which appears to behave as a strong loss-of-function model given the absence of protein, reduced body weight and neonatal lethality at the late embryonic and early postnatal stages.

### *TTC5^R263*/R263*^* mutation causes reduced brain size and abnormal cortical development

Evaluation of the gross brain morphology of *TTC5^R263*/R263*^* mice at P0 showed significant decreases in total brain area (mutant 27.70 ± 2.26mm^2^ vs. control 30.79 ± 1.13mm^2^), cortical area (mutant 17.15 ± 1.65mm^2^ vs. control 19.38 ± 0.81mm^2^), anteroposterior length (mutant 4.13 ± 0.21mm vs. control 4.46 ± 0.11mm), and cortical length (mutant 4.26 ± 0.20mm vs. control 4.64 ± 0.11mm) (**Fig. 2a-d**). These reductions in brain dimensions occur alongside the decreased body weight observed in *TTC5^R263*/R263*^* pups (**Fig. 1g, h**), indicating that reduced neonatal brain size occurs in the context of broader growth impairment.

**Fig. 2.**
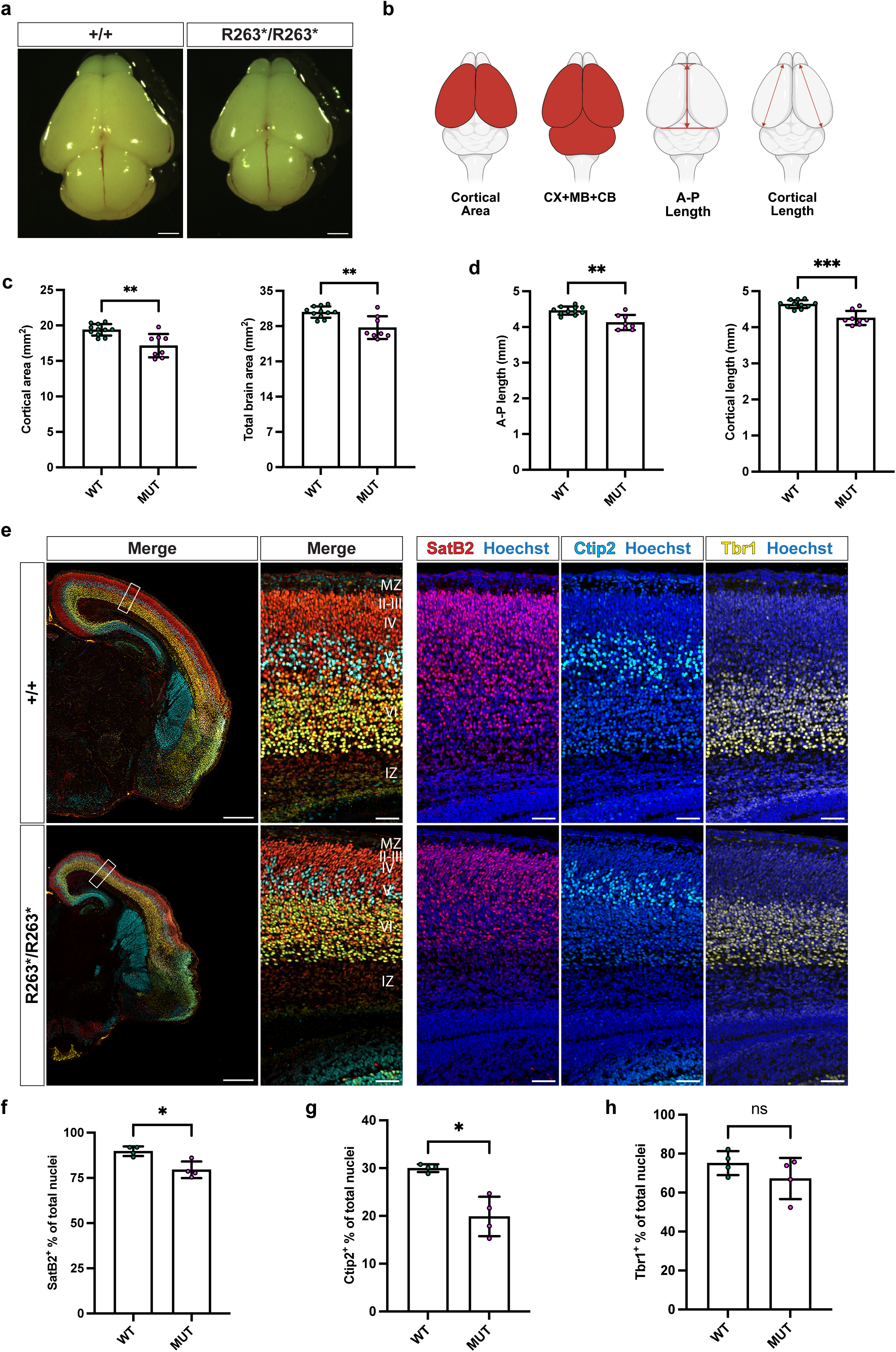
Decreased brain size and reduced cortical neurons in neonatal *TTC5^R263*/R263*^* mice. **a** Representative images of whole brains of wild-type (left) and *TTC5^R263*/R263*^* (right) mice at P0. Scale bars, 1 mm. **c**, **d** Measurements of brain size and cortical parameters were taken at P0 (as illustrated in **b**), including the cortex (CX), the midbrain (MB), the cerebellum (CB), the anteroposterior length of the cortex (A-P length), and the average length of the two cortical hemispheres (cortical length). At P0, *TTC5^R263*/R263*^* brains show decreased cortical area (**, *p* = 0.006), total brain area (**, *p* = 0.0055), A-P length (**, *p* = 0.0024), and cortical length (***, *p* = 0.0005) compared with wild-type littermates. n = 11 WT (6 males and 5 females) and 8 MUT (6 males and 2 females) animals from 4 litters. **e** Representative images of corresponding coronal brain sections from mid-posterior cortex of P0 wild-type and *TTC5^R263*/R263*^*brains immunostained with cortical neuronal markers Satb2, layers II-IV (red); Ctip2, layer V (cyan); and Tbr1, layer VI (yellow). Sections were counterstained with Hoechst 33342 (blue) to label nuclei. Merged images are marked with cortical layers II-VI along with the marginal zone (MZ) and the intermediate zone (IZ). Scale bars, 500*μ*m, 50*μ*m. **f-h** Quantification of cortical layer marker-positive cells in 200*μ*m wide fields taken from the primary visual cortex (V1). *TTC5^R263*/R263*^* mutants exhibit significant reduction in neuronal populations in the upper layers of the cortical plate: percentage of Satb2^+^ neurons to total nuclei (*, *p =* 0.013) and percentage of Ctip2^+^ neurons to total nuclei (*, *p* = 0.014). Deep layer Tbr1^+^ percentage in *TTC5^R263*/R263*^*brains is mildly decreased in comparison to wild type but is not statistically significant (*p =* 0.25). n = 4 WT (2 males and 2 females) and 4 MUT (2 males and 2 females) animals from 4 litters. Data are presented as means ± SDs. Statistical analyses were conducted using unpaired two-tailed t test with Welch’s correction.

To further examine the development of the neocortex and evaluate the cellular composition of the postnatal cortical plate (CP), we performed immunohistochemistry on newborn TTC5 mouse brains using layer-specific cortical neuronal markers Tbr1(layer VI)^52^,^53^, Ctip2 (layer V)^54^, and Satb2 (layer II-IV)^55,56^. *TTC5^R263*/R263*^* neocortex displayed correct positioning of cortical neurons and relatively preserved layering of the cortical plate (**Fig. 2e**). Despite the apparent overall normal patterning, there is thinning of the cerebral cortex, with the most profound reduction in the superficial cortical layers. Neuronal populations were significantly decreased in *TTC5^R263*/R263*^*mutants in comparison with the controls in layers II-IV, as illustrated by the ratio of Satb2^+^ neurons to total layer II-IV nuclei (mutant 79.48 ± 4.60% vs. control 89.76 ± 2.64%), and in layer V, by the ratio of Ctip2^+^ neurons to total layer V nuclei (mutant 19.88 ± 4.13% vs. control 30.00 ± 0.82%) (**Fig. 2f, g**). Tbr1^+^ neurons in layer VI of the *TTC5^R263*/R263*^* CP exhibit a more modest and nonsignificant decrease (mutant 67.21 ± 10.58% vs. control 75.13 ± 6.16%) (**Fig. 2h**). Therefore, neonatal *TTC5^R263*/R263*^* animals have fewer total neurons and reduction across cortical layers with a gradient in severity, with most severe reductions in the most superficial layers and least severe in the deepest layers. As the cortical layers are formed in an inside-out fashion, where the deepest layers are composed of early-born neurons with sequential migration of later-born neurons towards the upper layers, this process is tightly regulated by temporal identities of neuronal precursors throughout different developmental stages of the neocortex.^57–59^ Our data suggest that germline TTC5 deficiency results in early and lasting defects in neurogenesis with more profound reduction in later-born superficial neurons.

### Proliferative defects in *TTC5^R263*/R263*^* embryonic cortical neuron progenitors

Our observations in newborn animals prompted us to further investigate the effects of TTC5 deficiency on neurogenesis during embryonic development. Crown-Rump length (CRL) measurements were used to confirm the developmental age of E14.5 embryos.^60^ No significant difference in CRL was observed among the different genotypes (WT: 12.28mm ± 0.8298; HET 13.09mm ± 1.115; MUT: 13.07mm ± 1.183) (**Supplementary Fig. 3**). At E14.5, during mid-neurogenesis, we observed decreases in overall cortical thickness (**Fig. 3a**). Immunostaining of doublecortin (Dcx), a marker for migrating immature neurons^61^, showed thinning of both the neuronal layer (Dcx^+^, CP and IZ) (mutant 202.9 ± 13.12 *μ*m vs. control 252.7 ± 19.92 *μ*m) and the neurogenic layer (Dcx^-^, VZ and SVZ) (mutant 91.83 ± 1.44 *μ*m vs. control 169.5 ± 19.34 *μ*m) of the *TTC5^R263*/R263*^* developing cortex in comparison to wild type (**Fig. 3b**). Despite the reduction in both regions, *TTC5^R263*/R263*^* embryos had a higher ratio of Dcx^+^ layer to the total cortical thickness (mutant 68.8 ± 1.33% vs. control 59.9 ± 1.70%), suggesting that a higher proportion of the *TTC5^R263*/R263*^*cortex is composed of differentiating neurons at E14.5, with a more dramatic reduction in the neurogenic region populated by progenitor cells (**Fig. 3c**). At this developmental stage, the deepest layer (layer VI) of the cortical plate is being generated and Tbr1 is expressed homogeneously in the superficial layer of the neocortex.^52^ Tbr1 immunostaining at E14.5 showed fewer early-born postmitotic neurons in the *TTC5^R263*/R263*^* CP (mutant 130.7 ± 23.26 vs. control 192.1 ± 14.5) (**Fig. 3d, e**).

**Figure 3.**
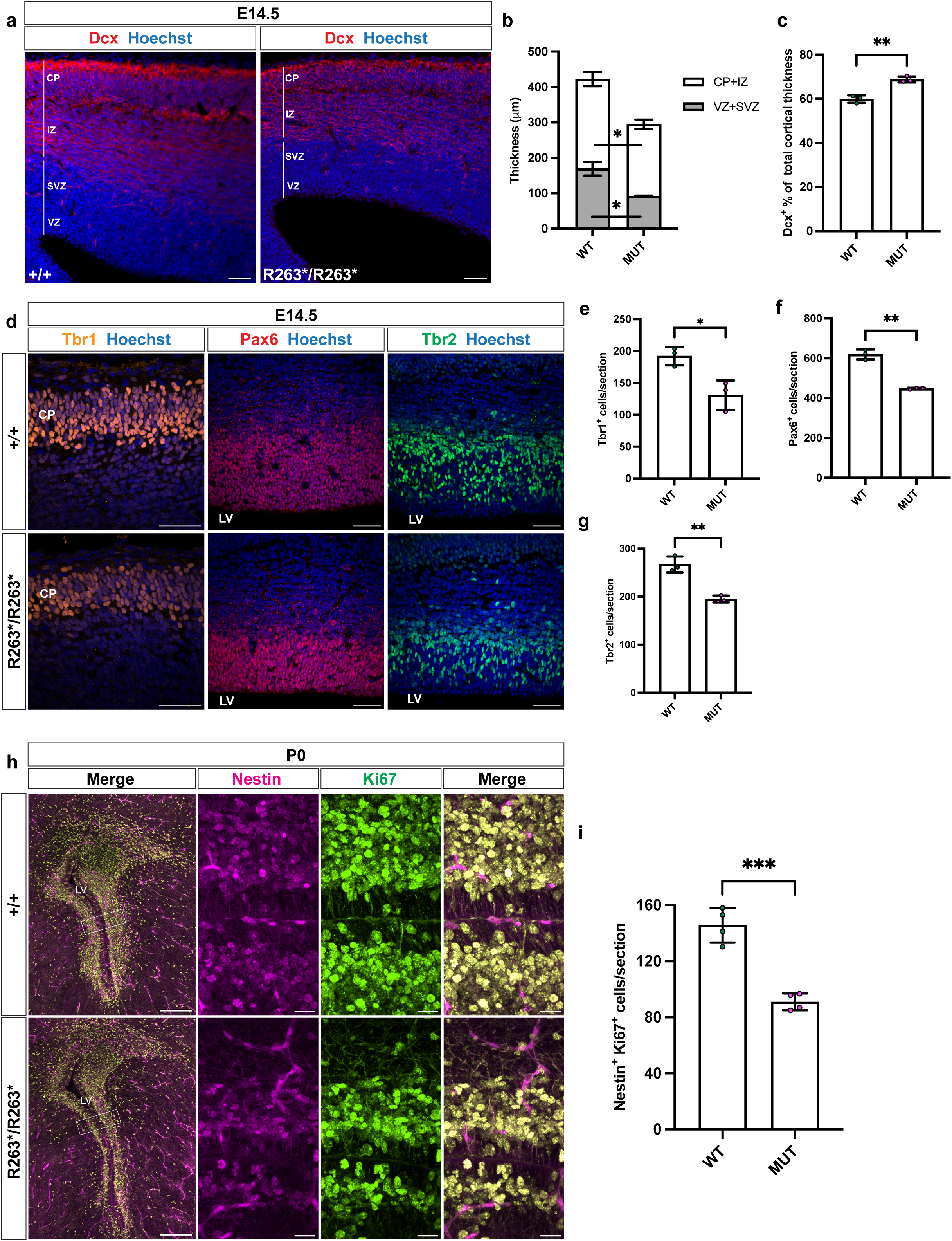
Impaired neurogenesis and loss of progenitors in *TTC5^R263*/R263*^* mice. **a** Dcx (red) immunostaining of coronal sections in the mid-anterior cortex of E14.5 wild-type and *TTC5^R263*/R263*^* brains. Dcx, a marker for immature neurons, shows the neuronal layer of the cortex, including the cortical plate and the intermediate zone. Scale bars, 50*μ*m. CP, cortical plate; IZ, intermediate zone; SVZ, subventricular zone; VZ, ventricular zone. **b** Quantification of the thickness of the Dcx^+^ neuronal layer (CP and IZ) and the Dcx^-^neurogenic layer (VZ and SVZ). *TTC5^R263*/R263*^* animals show reduction in cortical thickness in both the neuronal layer (*, *p =* 0.029) and the neurogenic layer (*, *p =* 0.02). **c** A higher fraction of the *TTC5^R263*/R263*^*developing cortex is neuronal at E14.5 despite overall reduction of cortical thickness demonstrated as the ratio of Dcx-positive cortical thickness over total cortical thickness (**, *p =* 0.0025). **d** At E14.5, the number of early-born postmitotic neurons was analyzed by immunostaining of Tbr1 (orange) while neuronal progenitors were analyzed by Pax6 (radial glial cells, red) and Tbr2 (intermediate progenitors, green) staining. Nuclei are stained with Hoechst 33342 (blue). Scale bars, 50*μ*m. LV, lateral ventricle. **e** Quantification of Tbr1^+^ neurons in the CP of E14.5 littermates. *TTC5^R263*/R263*^* embryos show reduced neuronal number in comparison to the wild type (*, *p =* 0.025). **f, g** Quantification of Pax6^+^ radial glia and Tbr2^+^ intermediate progenitors in the VZ and SVZ. At E14.5, both populations of neuronal precursors are reduced in the *TTC5^R263*/R263*^* embryos: Pax6^+^ progenitors (**, *p =* 0.0057) and Tbr2^+^ progenitors (**, *p =* 0.0086). n = 3 WT (1 male and 2 females) and 3 MUT (1 male and 2 females) animals from 2 litters. **h** Coronal sections immunostained with nestin (magenta), a marker for NPCs, co-labeled with cell proliferation marker, Ki67 (green), in P0 wild-type and *TTC5^R263*/R263*^* brains. Images were taken from the neuroepithelium (ne) around the lateral ventricle (LV) of anterior cortex. Scale bars, 200*μ*m, 20*μ*m. **i** Quantification of proliferating neural progenitor cells by assessing the total number of co-labeled nestin^+^ Ki67^+^ cells in a 200*μ*m × 200*μ*m area in the neuroepithelium. *TTC5^R263*/R263*^* mice show a decrease in the number of co-labeled nestin^+^/Ki67^+^ cells (***, *p =* 0.0009). n = 4 WT (2 males and 2 females) and 4 MUT (2 males and 2 females) animals from 4 litters. Data are presented as means ± SDs. Statistical analyses were conducted using unpaired two-tailed t test with Welch’s correction.

During embryonic development, the spatial-temporal regulation of cortical neuronal progenitors is established by the sequential expression of transcription factors: Pax6 in radial glia (RGCs), Tbr2 in intermediate progenitors (IPs), and Tbr1 in postmitotic neurons.^62^ Immunostaining of Pax6 and Tbr2 at E14.5 showed loss of both RGCs (mutant 448.1 ± 4.08 vs. control 619.7 ± 24.55) and IPs (mutant 195.1 ± 6.99 vs. control 267.2 ± 16.54) in the ventricular zone (VZ) and subventricular zone (SVZ) of the *TTC5^R263*/R263*^* developing cortex compared to wild type (**Fig. 3f, g**). Together, our data suggest a premature depletion of the neuronal progenitor pool in both sub-populations during early stages of cortical neurogenesis caused by the *TTC5^R263*/R263*^* mutation, likely leading to observed deficits across the CP in postnatal animals.

While postmitotic neurons in the cerebral cortex are generated predominantly during embryogenesis, neural progenitor cells (NPCs) remain in the neurogenic regions in the VZ and SVZ around the lateral ventricle into the early postnatal period.^63,64^ During later stages of neurogenesis in neonatal brains, NPCs are less committed to neuronal lineage through an NPC fate switch to astrogliogenesis by repressing *Ngn1* expression.^63,65,66^ Here, by using an anti-nestin marker, an intermediate filament protein that is exclusively expressed in uncommitted NPCs, we further examined continued neural development in the neonatal cortex of *TTC5^R263*/R263*^* mutants.^64,67,68^ Double immunostaining of nestin and proliferation marker, Ki67 revealed fewer dividing NPCs in the VZ and SVZ (**Fig. 3h**) of P0 *TTC5^R263*/R263*^* brains compared to wild type. Co-labeled nestin^+^/Ki67^+^ cells were decreased by almost 40% in the *TTC5^R263*/R263*^* neuroepithelium compared to wild type (mutant 91.06 ± 6.01 vs. control 145.7 ± 12.36) (**Fig. 3i**). This reduction in NPC population in neonatal animals further suggests that TTC5 deficiency leads to long term defects in cortical progenitor maintenance that persists in postnatal mouse brains.

To understand the mechanism underlying the reduction of neuronal precursor populations in *TTC5^R263*/R263*^* embryos, we analyzed progenitor cell divisions in the neurogenic region of the E14.5 cortex. TTC5 depletion in HeLa cells has been demonstrated to lead to subtle but reproducible increase in mitotic duration.^23,25^ We therefore assessed mitotic-stage distributions of RGCs by the localization of mitotic marker phospho-Histone H3 (pH3) and mitotic spindle marker aurora kinase B (AURKB) (**Supplementary Fig. 4**). A significantly higher fraction of *TTC5^R263*/R263*^*RGCs were found in prometaphase and metaphase compared with controls (mutant 64.30 ± 2.07% vs. control 38.87 ± 4.46%) (**Fig. 4a, b**). *TTC5^R263*/R263*^* RGCs also exhibited increased mitotic index by the percentage of pH3-positive cells out of total Pax6-positive cells (mutant 10.76 ± 1.27% vs. control 6.22 ± 0.62%) (**Fig. 4c, d**). The aberrant loss of the progenitor pool, together with the altered mitotic progression profile, suggest a prometaphase and/or metaphase delay/arrest in the *TTC5^R263*/R263*^* RGCs. Given the marked reduction in tubulin abundance and impaired microtubule dynamics observed in *TTC5^R263*/R263*^*brains (**Fig. 6-7**), these findings are consistent with a potential involvement of microtubule dysfunction in the observed mitotic abnormalities.^69–72^

**Figure 4.**
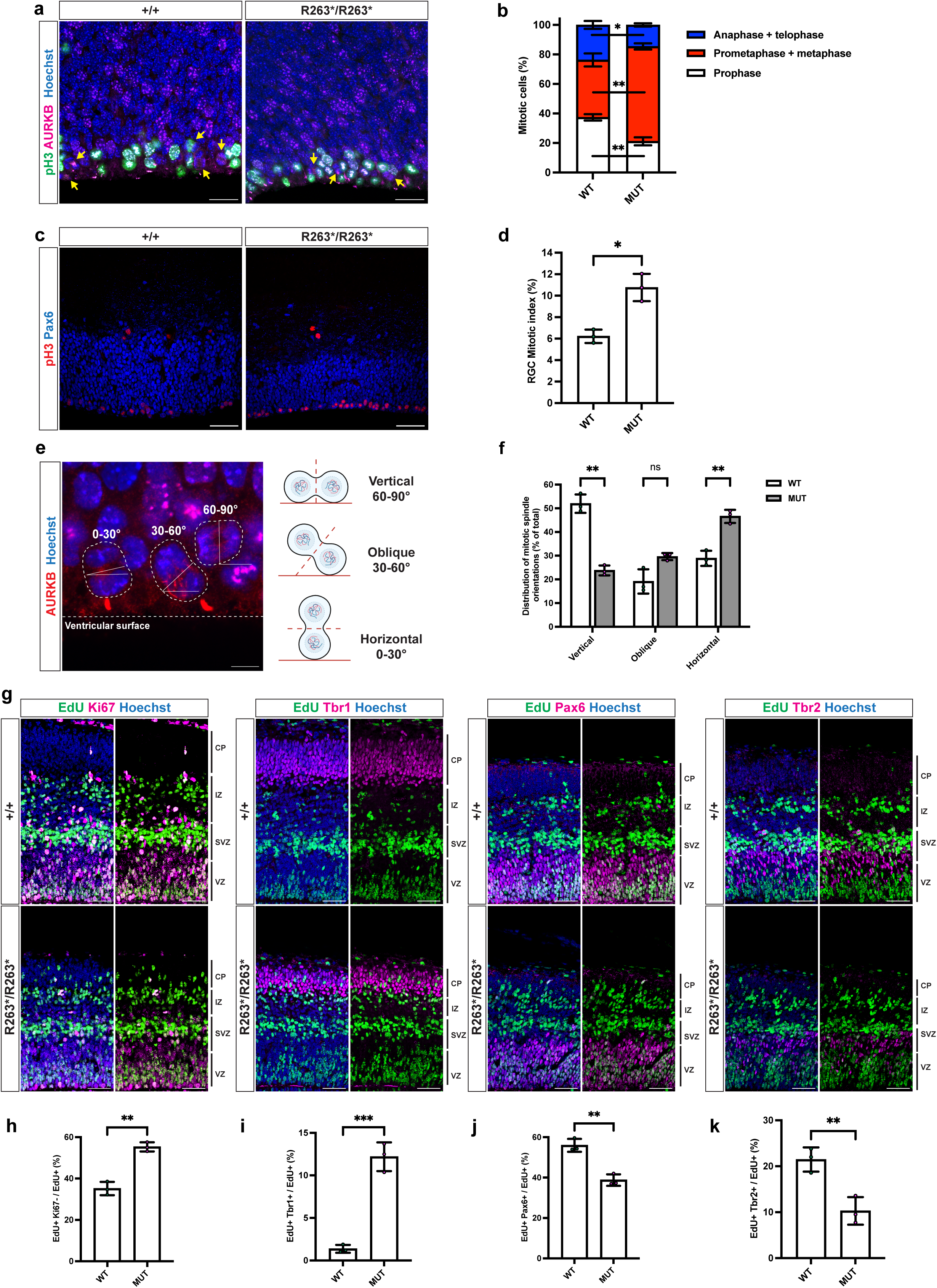
*TTC5^R263*/R263*^* RGCs have mitotic defects and progenitor dynamics. **a** Representative images of M-phase RGCs at the ventricular surface at E14.5. Yellow arrows depict RGCs from anaphase to telophase. Scale bars, 20*μ*m. **b** Analysis of mitotic progression profile of *TTC5^R263*/R263*^* cortical progenitors at E14.5. The percentage of progenitor cells in the VZ of *TTC5^R263*/R263*^*developing cortex in prometaphase to metaphase (**, *p =* 0.0037) is significantly increased, while fractions of prophase (**, *p =* 0.0015) and anaphase to telophase (*, *p =* 0.017) cells are significantly decreased compared to wild type. An average of 325 WT and 388 MUT cells were counted/animal. **c,d** Analysis of RGC mitotic index as the percentage of pH3^+^/Pax6^+^ double positive cells to the total number of Pax6^+^ cells. The mitotic index of *TTC5^R263*/R263*^* RGCs is significantly increased compared to control (*, *p =* 0.013). Scale bars, 50*μ*m. **e, f** Analysis of mitotic spindle orientation in E14.5 cortical progenitors. Anaphase and telophase progenitor cells are classified into three types based on the angle of the cleavage plane relative to the ventricular surface (dotted line): horizontal, 0-30°; oblique, 30°-60°; vertical, 60°-90°. Central spindle and midbody are visualized by AURKB staining (red) and nuclei are visualized by Hoechst 33342 (blue). The distribution of the three classes of mitotic progenitors is altered in the *TTC5^R263*/R263*^* brains. *TTC5^R263*/R263*^* progenitor cells have decreased vertical (**, *p =* 0.0015) and increased horizontal (**, *p =* 0.0021) and oblique (ns, *p =* 0.062) cleavage planes in comparison to the wild type. An average of 77 WT and 57 MUT cells were counted/animal. Scale bars: 5*μ*m. **g** E14.5 brain sections 24 hours after EdU injection and stained for EdU, Ki67, Tbr1, Pax6, and Tbr2. Scale bars: 50*μ*m. **h-k** Quantification of the fractions of EdU-labeled cells that were Ki67-negative (**, *p =* 0.0015), Tbr1-positive (***, *p =* 0.0004), Pax6-positive (**, *p =* 0.0022), and Tbr2-positive (**, *p =* 0.0084). n = 3 WT (1 male and 2 females) and 3 MUT (1 male and 2 females) animals from 2 litters. Data are presented as means ± SDs. Statistical analyses were conducted using unpaired two-tailed t test with Welch’s correction.

**Figure 5.**
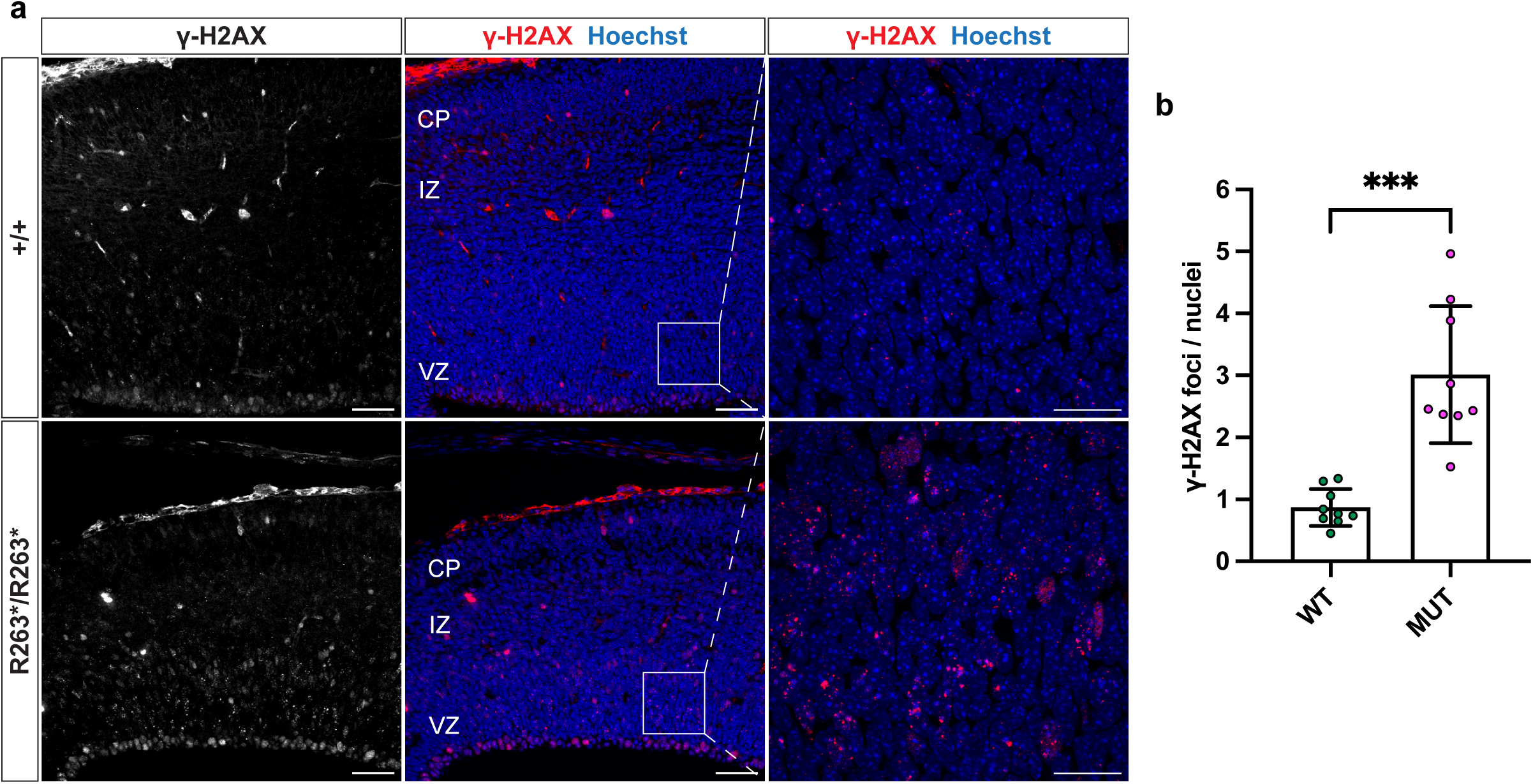
Elevated DNA damage in *TTC5^R263*/R263*^* neuronal progenitors. **a** At E14.5, the level of DNA damage in NPCs is analyzed by immunostaining of γ-H2AX (red). Rectanglar areas are shown on the right with higher magnification. Nuclei are counterstained with Hoechst 33342 (blue). Scale bars, 50*μ*m, 20*μ*m. **b** Quantification of the number of nuclear γ-H2AX foci in the ventricular zone. *TTC5^R263*/R263*^* progenitors show increased number of γ-H2AX foci per nucleus (***, *p =* 0.0003). n = 3 WT (1 male and 2 females) and 3 MUT (1 male and 2 females) animals from 2 litters. Data are presented as means ± SDs. Statistical analyses were conducted using unpaired two-tailed t test with Welch’s correction.

**Fig. 6.**
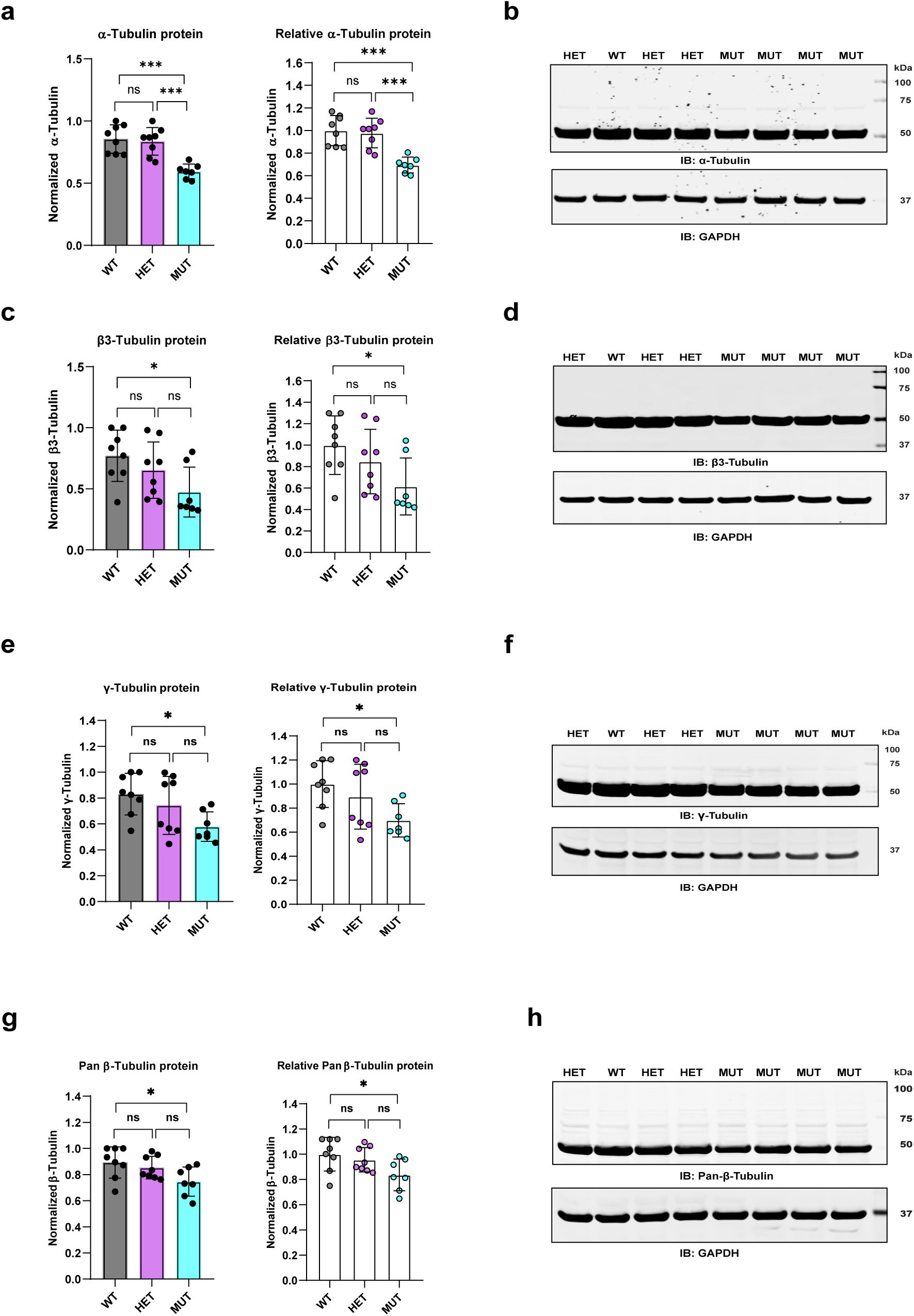
Reduced tubulin protein expression in *TTC5^R263*/R263*^* mutant brains. **a,b** Western blotting was performed to determine α-tubulin protein expression in *TTC5^R263*/R263*^*mouse brains. Representative gel images are shown in **b**. α-tubulin protein was normalized to GAPDH and compared between *TTC5^+/+^*, *TTC5^+/R263*^*, and TTC5*^R263*/R263*^* animals (***, *p* = 0.0002 for WT vs. MUT; ***, *p* = 0.0004 for HET vs. MUT; ns, p=0.9242 for WT vs. HET). One-way Anova with Tukey’s multiple comparisons test. **c, d** Quantification and comparison of β3-tubulin between *TTC5^+/+^*, *TTC5^+/R263*^*, and TTC5*^R263*/R263*^* animals (*, *p* = 0.0391 for WT vs. MUT; ns, *p* = 0.2682 for HET vs. MUT; ns, p=0.5337 for WT vs. HET). One-way Anova with Tukey’s multiple comparisons test. Representative gel images are shown in **d**. **e, f** Quantification and comparison of γ-tubulin between *TTC5^+/+^*, *TTC5^+/R263*^*, and TTC5*^R263*/R263*^* animals (*, *p* = 0.0303 for WT vs. MUT; ns, *p* = 0.1914 for HET vs. MUT; ns, p=0.5891 for WT vs. HET). One-way Anova with Tukey’s multiple comparisons test. Representative gel images are shown in **f**. **h, i** Quantification and comparison of pan-β-tubulin between *TTC5^+/+^*, *TTC5^+/R263*^*, and TTC5*^R263*/R263*^* animals (*, *p* = 0.0387 for WT vs. MUT; ns, *p* = 0.1520 for HET vs. MUT; ns, p=0.7465 for WT vs. HET). One-way Anova with Tukey’s multiple comparisons test. Representative gel images are shown in **i.** n = 8 WT (3 males and 5 females), 8 HET (6 males and 2 females) and 7 MUT (2 males and 5 females) animals.

**Fig. 7.**
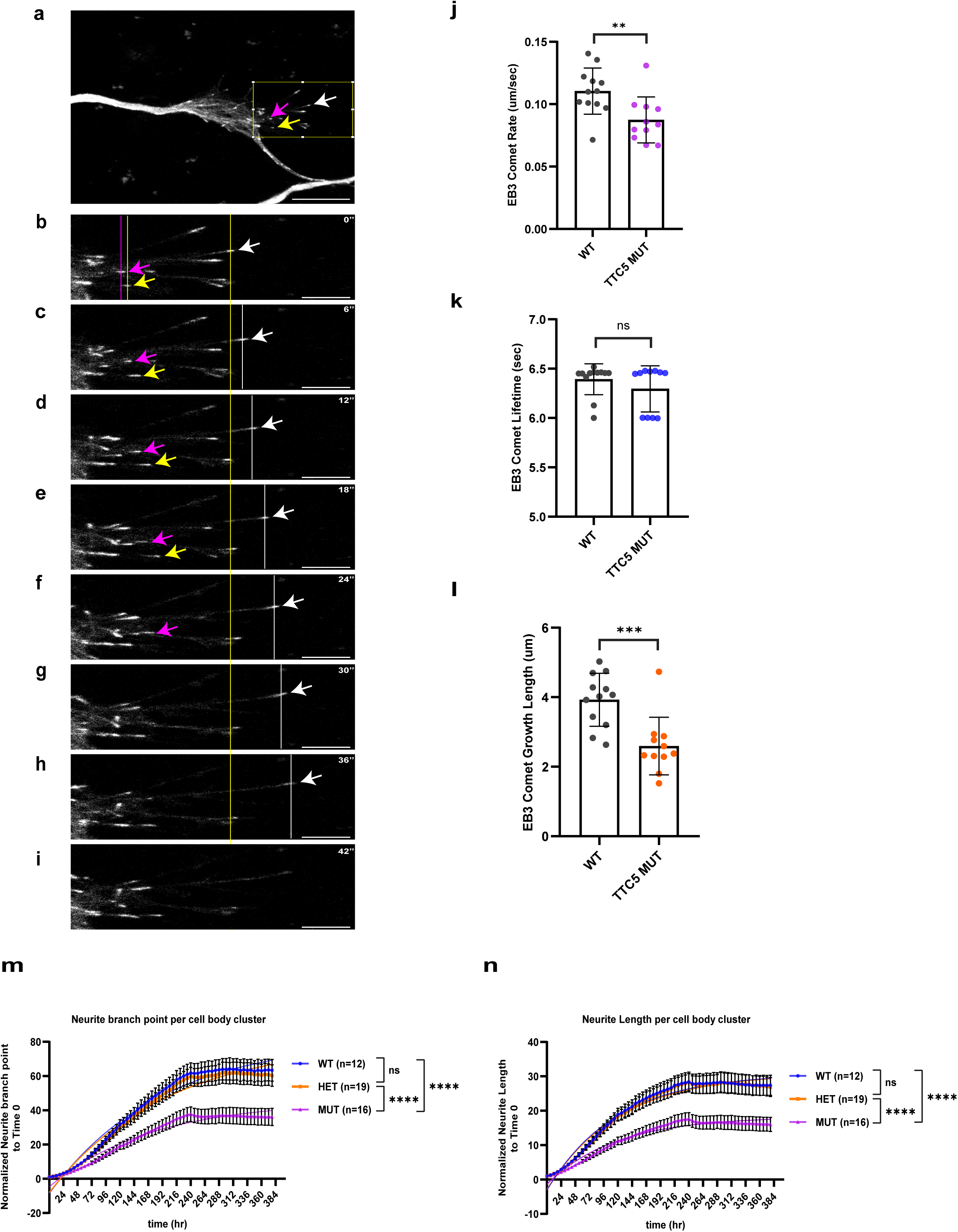
*TTC5^R263*/R263*^* affects MT behavior in cortical neurons and reduces neuronal arborization. Cortical neurons were transfected with EB3-tdTomato at DIV2, and live cells were analyzed on DIV3 by confocal microscopy in a 37°C chamber. EB3-tdTomato comet movements were imaged and recorded. The rectangles in (**a**) are present the areas magnified in (**b-i**). The images in **b-i** were acquired every 6 sec. Arrows in these panels indicate the beginning and end positions of a selected number of EB3-tdTomato comets. Different colored arrows help to distinguish the movement of different comets. Scale bars: A, 10 *μ*m; B, 5 *μ*m. EB3 comet rate (*μ*m/sec), lifetime (sec) and growth length (*μ*m) were measured by Image J. Data were collected from 106 comets from 24 WT neurons and 141 comets from 36 MUT neurons from 6 independent experiments. Statistics were analyzed at the animal-level by unpaired two-tailed t test with Welch’s correction for comparisons between WT and MUT: EB3 comet rate (**, *p =* 0.0068) (**j**); EB3 comet lifetime (ns, *p =* 0.26) (**k**) and EB3 comet growth length (***, *p =* 0.0007) (**l**). n = 12 WT (8 males and 4 females) and 11 MUT (5 males and 6 females) animals from 10 litters. **m, n** Measurement of neurite outgrowth with respect to branch points (**m**) and neurite length (**n**) in cortical neurons from TTC5*^R263*/R263*^*mouse line. Nonlinear fit comparison was performed for neurite branch points (****, *p* < 0.0001 for WT vs. MUT; ****, *p* < 0.0001 for HET vs. MUT; ns, *p* = 0.1241 for WT vs. HET) and for neurite length (****, *p* < 0.0001 for WT vs. MUT; ****, *p* < 0.0001 for HET vs. MUT; ns, *p* = 0.9321 for WT vs. HET). n = 12 WT (8 males and 4 females), 19 HET (9 males and 10 females) and 16 MUT (8 males and 8 females) animals from 10 litters. Data are presented as means ± SDs.

Neural progenitor cell fate determination has been proposed to be influenced by the orientation of the mitotic division plane relative to the ventricular surface.^73–75^ Vertical cleavage planes (60°-90°) are more commonly associated with symmetric cell divisions, whereas horizontal (0-30°) or oblique (30°-60°) cleavage planes are associated with asymmetric cell fates.^73–75^ During cortical development, RGCs predominantly divide with cleavage planes perpendicular to the ventricular surface to produce side-by-side daughter cells, and alterations in mitotic orientation contribute to abnormal progenitor behavior.^76,77^ *TTC5^R263*/R263*^* RGCs showed decreased cell divisions with vertical cleavage planes (mutant 23.79 ± 2.08% vs. control 51.97 ± 3.89%) and increased horizontal (mutant 46.59 ± 2.78% vs. control 28.90 ± 3.19%) and oblique (mutant 29.62 ± 1.48% vs. control 19.13 ± 5.13%) cleavage planes (**Fig. 4e, f**). To further assess cell cycle exit and progeny fate in *TTC5^R263*/R263*^* embryos, we labeled progenitors with EdU at E13.5 and examined cell-cycle exit using EdU/Ki67 double labeling and quantified EdU-positive cells expressing neuronal (Tbr1), radial glial (Pax6), and intermediate progenitor (Tbr2) markers at E14.5 **(Fig. 4g**). Cell cycle exit was increased by 1.6-fold in *TTC5^R263*/R263*^* animals by the ratio of EdU-positive Ki67-negative cells to the total number of EdU-positive cells (mutant 55.30 ± 2.24% vs. control 35.17 ± 3.23%) (**Fig. 4h**). Despite the already apparent thinning of the CP at E14.5, the EdU^+^ population in the *TTC5^R263*/R263*^* embryos resulted in increased proportion of early-born postmitotic neurons (mutant 12.19 ± 1.69% vs. control 1.37 ± 0.46%), concomitant with reduction in both RGCs (mutant 38.78 ± 2.84% vs. control 55.99 ± 3.19%) and IPs (mutant 10.28 ± 3.00% vs. control 21.47 ± 2.64%) (**Fig. 4i-k**). The observed alterations in progeny composition suggest a shift toward neuronal differentiation at the expense of progenitor maintenance. In summary, these data demonstrate a premature depletion of neuronal progenitors during mid-early neurogenesis associated with mitotic defects and cell fate alterations in the *TTC5^R263*/R263*^* developing cortex.

TTC5*-*depleted HeLa cells show evidence of erroneous mitosis^23–25^, which has been associated with DNA damage generation.^78–80^ To assess DNA damage signaling in *TTC5^R263*/R263*^* NPCs, we examined phospho-histone H2AX (γ-H2AX), a marker for cellular DNA damage response (**Fig. 5a**). We observed significantly elevated nuclear γ-H2AX foci (mutant 3.01 ± 1.11 vs. control 0.87 ± 0.30) (**Fig. 5b**), indicating accumulation of DNA damage. We further investigated whether the mitotic defects and increased DNA damage led to increased apoptosis, resulting in the loss of neuronal and/or progenitor cell populations in the *TTC5^R263*/R263*^* developing cortex. For cell death analysis, brain sections were examined using activated cleaved caspase 3 (CC3) immunostaining and terminal deoxyribonucleotidyl transferase mediated dUTP-biotin nick end labeling (TUNEL) assay, both markers for cells undergoing apoptosis. No elevation in apoptosis was detected in the *TTC5^R263*/R263*^* cortical plate and neuroepithelium at P0 and E14.5, and signal patterns of the markers across the control and mutant cortices resemble naturally occurring cell death in development (**Supplementary Fig. 5, 6**).^81–83^ Therefore, these findings support impaired expansion capacity of *TTC5^R263*/R263*^* NPCs as the major contributing mechanism underlying the observed cortical deficits.

### Transcriptome analysis in neonatal *TTC5^R263*/R263*^* cortex demonstrates altered molecular pathways, including dysregulation of tubulin mRNA abundance and cell cycle genes

To study the function of TTC5 at the whole transcriptome level, we performed bulk RNA-sequencing on total mRNA from P0 mouse cortex. To explore the biological variations between wild type and *TTC5^R263*/R263*^* animals, we first performed a principal component analysis (PCA) on samples across all 16375 expressed genes (**Supplementary Fig. 7a; Supplementary Table 1**). PC1 explained 40.4% of variance across all samples and can separate wild-type and *TTC5^R263*/R263*^* samples into two well-separated clusters, indicating a distinct transcriptome profile of mutants compared with wild type.

Following a standard bulk RNA-seq analysis pipeline (STAR-StringTie-DESeq2), 592 genes were identified as significant differentially expressed genes (DEGs, absolute log2FC greater than 1 with p.adj less than 0.05, **Supplementary Table 2**), including 240 upregulated genes and 352 downregulated genes. Heatmaps of the top 100 DEGs are presented in **Supplementary Fig. 7b**. In addition to this DEG set, we separately examined cell-cycle regulators and tubulin-related genes based on their established functions and biological relevance. *TTC5* expression was downregulated in the *TTC5^R263*/R263*^* cortex with log2FC of - 0.855 and p.adj of 6.39e-11.

We observed dysregulated genes that may be associated with the mitotic defects described above. Prolonged mitosis, particularly prolonged prometaphase, has been associated with altered cell fate and blocked cell proliferation in producing G1-arrested progeny.^84,85^ Transcriptional profile of the *TTC5^R263*/R263*^* developing cortex revealed altered expression of cell cycle progression mediating cyclin-dependent kinases (CDKs), their regulatory cyclins, and CDK inhibitors (**Supplementary Table 3**). In particular, mRNA levels were downregulated in *TTC5^R263*/R263*^*cortices for CDK4 (by 25%, *padj* = 0.0029) and its associated D-type cyclins (cyclin D1 by 24%, *padj* = 0.056, ns; cyclin D2 by 42%, *padj* = 7.01×10^-10^), as well as CDK2 (by 18%, *padj* = 0.035) and its associated E-type cyclins (cyclin E1, no change; cyclin E2 by 28%, *padj* = 0.014). Conversely, expression of INK4 family proteins p16^INK4A^ (CDKN2A, by 2.3-fold, *padj* = 0.02) and p15^INK4B^ (CDKN2B, by 2-fold, *padj* = 9.2×10^-4^), and Cip/Kip family proteins p21^Cip1/Waf1^ (CDKN1A, by 2.3-fold, *padj* = 3.45×10^-5^) and p57^Kip2^ (CDKN1C, by 2.3-fold, *padj* = 1.84×10^-4^) was upregulated.

To further examine cell-type-specific changes in cell-cycle associated gene expression, we performed pseudobulk differential expression analysis using our single nuclei RNA sequencing dataset. We observed nominal expression differences (*p < 0.05*) in several G1/S cell-cycle regulators, including decreased CDK4 expression in Cycling Dlx+ inhibitory progenitors (by 49%) and Cycling radial glia/glial progenitors (by 60.7%), cyclin D2 expression in Cycling Dlx+ inhibitory progenitors (by 29.9%) and Early differentiating neurons (by 26.6%), and CDK6 expression in Cycling radial glia/glial progenitors (by 19%). CDKN1A expression was increased by 6.1-fold in Cycling radial glia/glial progenitors. Together, these findings indicate altered expression of cell-cycle-associated genes in the mutant cortex, consistent with the observed abnormalities in progenitor populations and proliferative behavior.

TTC5 has been proposed as a tubulin-specific ribosome-associating factor that regulates degradation of tubulin mRNAs in response to excess soluble tubulin.^23^ Therefore, we examined tubulin gene expression in the bulk RNA sequencing dataset. A total of 24 mouse tubulin genes were identified: Tub, Tuba1a, Tuba1b, Tuba1c, Tuba4a, Tuba8, Tubb1, Tubb2a, Tubb2a-ps2, Tubb2b, Tubb3, Tubb4a, Tubb4b, Tubb5, Tubb6, Tubd1, Tube1, Tubg1, Tubg2, Tubgcp2, Tubgcp3, Tubgcp4, Tubgcp5, Tubgcp6 (**Table 1**; **Supplementary Table 2, 4**). Among them, 11 genes (orange) were significantly downregulated. Interestingly, Tubb3, a major tubulin subtype in neurons, was significantly downregulated (by 31%, *p.adj* = 2.15x 10^-5^). Two genes (blue) were significantly upregulated. The remaining 11 tubulin genes were not significantly changed. We further explored pathways affected by the TTC5 dysfunction through pathway enrichment analysis on DEGs (**Supplementary Table 5)**. G protein-coupled receptors (GPCRs) were significantly enriched among DEGs (p.adj = 0.0238).

**Table 1.** Tubulin genes from Bulk RNA sequencing

|  | **log2FoldChange** | **pvalue** | **padj** | **expression<br>relative to wt** |
|---|---|---|---|---|
| Tub | 0.176081118 | 0.02844912 | 0.06942541 | 1.130 |
| Tuba1a | -0.161975826 | 0.06814497 | 0.14114334 | 0.894 |
| **Tuba1b** | -0.409370529 | 8.90E-05 | **0.00053641** | **0.753** |
| Tuba1c | -0.159702985 | 0.07236372 | 0.1483413 | 0.895 |
| Tuba4a | -0.266536718 | 0.08748127 | 0.17203479 | 0.831 |
| Tuba8 | 0.224673627 | 0.62331113 | 0.74590406 | 1.165 |
| Tubb1 | 0.032481526 | 0.96659283 | 0.98018627 | 1.021 |
| **Tubb2a** | -0.369696631 | 2.17E-07 | **3.36E-06** | **0.774** |
| **Tubb2a-ps2** | -0.406125976 | 0.00972474 | **0.02870908** | **0.755** |
| **Tubb2b** | -0.29789446 | 0.00120701 | **0.00498554** | **0.813** |
| **Tubb3** | -0.535415292 | 1.93E-06 | **2.15E-05** | **0.690** |
| **Tubb4a** | -0.388979109 | 7.57E-05 | **0.00046927** | **0.764** |
| **Tubb4b** | -0.404185545 | 3.36E-06 | **3.42E-05** | **0.756** |
| Tubb5 | -0.206060757 | 0.02421791 | 0.06089258 | 0.867 |
| **Tubb6** | 0.520085462 | 0.01853204 | **0.04890804** | **1.434** |
| **Tubd1** | -0.946209312 | 0.00600997 | **0.01927804** | **0.519** |
| Tube1 | -0.200325823 | 0.17198833 | 0.29330839 | 0.870 |
| **Tubg1** | -0.302356035 | 2.95E-05 | **0.0002103** | **0.811** |
| **Tubg2** | 0.234603283 | 0.00085692 | **0.00372042** | **1.177** |
| Tubgcp2 | 0.211163707 | 0.02431524 | 0.06108359 | 1.158 |
| Tubgcp3 | 0.208192115 | 0.04123281 | 0.0939913 | 1.155 |
| **Tubgcp4** | -1.005499611 | 5.19E-06 | **4.93E-05** | **0.498** |
| Tubgcp5 | -0.170633519 | 0.03886573 | 0.08966579 | 0.888 |
| **Tubgcp6** | -0.482476908 | 1.12E-09 | **3.84E-08** | **0.716** |

We performed weighted gene co-expression network analysis (WGCNA) on *TTC5^R263*/R263*^* and wildtype samples . A scale-free co-expression network was constructed from 16,375 expressed genes, identifying 13 co-expression modules (**Supplementary Fig. 8a-b**, **9a**). A dendrogram showing network connection from genes is presented where each branch on the dendrogram represents a gene (**Supplementary Fig. 8b; Supplementary Table 6**). *Ttc5* was clustered in the brown module. Module correlation analysis revealed the strongest positive correlation between the brown and blue modules (**Supplementary Fig. 8c, 9a**), indicating that genes in these two modules may interact with the same biological processes. Pathway enrichment analysis on the brown module showed genes significantly associated with DNA binding transcription factor activity **(Supplementary Table 7)**. The blue module was enriched in genes associated with several neurological features and diseases, such as “abnormal communication”, “intellectual disability”, “neurodevelopmental abnormality”, and “abnormal brain morphology” (**Supplementary Table 7)**. Besides the blue module, the brown module also showed a high positive correlation with the green module. Importantly, the green module was significantly enriched with microtubule pathways (**Supplementary Table 7)**, including microtubule-based process, movement and microtubule bundle formation. The green module exhibited the second-highest positive correlation with Ttc5 expression (**Supplementary Fig. 8d, 9a**), further supporting an association between Ttc5 expression and microtubule-related gene networks.

The turquoise module presented the highest negative correlation with the brown module. Genes in the turquoise module was significantly enriched in ribosome biogenesis, nonsense-mediated mRNA decay, synapse translation, and mitotic progression and DNA damage checkpoints (**Supplementary Table 7)**. Given the large number of genes in the turquoise module (4,095), pathway enrichment analysis may identify broad biological associations that warrant cautious interpretation. In summary, bulk RNA sequencing of the neonatal *TTC5^R263*/R263*^*cortex revealed a distinct transcriptomic profile from wild-type controls, characterized by altered expression of tubulin-related and cell-cycle-associated genes. WGCNA further identified co-expression modules enriched in microtubule-related processes and neurodevelopmental pathways. Together, these findings demonstrate tissue-level transcriptional alterations consistent with the known biological functions of TTC5 and the observed cortical developmental phenotypes.

To further investigate the transcriptional alterations identified by bulk RNA-seq and examine gene expression changes within specific cortical cell populations, we performed single nuclei RNA sequencing (snRNAseq) of P0 wild-type and *TTC5^R263*/R263*^* cortices. We collected 174,444 single nuclei from wild-type (76,504; N=4) and *TTC5^R263*/R263*^* (97,940; N=4) mice. The overall number of nuclei was consistent across replicates **(Supplementary Fig. 10a)**. Using the integrated dataset of all animals, 16 distinct broad cell types were annotated using canonical markers **(Supplementary Fig. 10b)**. We subclassified excitatory neurons into cortical layer 1(1,074 nuclei, Reln and Trp73), layer 2-3 (15,625, Cux1 and Satb2), layer 4 (45,567, Rorb and Unc5d), layer 5-6 (18,430, Tbr1 and Foxp2) and immature excitatory (16,716, Dcx, Unc5d and Sema3c). We also identified Inhibitory neurons (53,273, Gad1 and Gad2), Neuronal progenitors (1,060, Eomes), Cycling Dlx+ inhibitory progenitors (2,145), Cycling radial glia/glial progenitors (2,253), Postmitotic neurons (2,129), Astrocytes (3,340, Aldh1l1), Microglia (1,231, Cx3cr1 and Csf1r), OPC and Oligodendrocytes (4,159, Pdgfra and Sox10), Endothelial (1,765, Pecam1 and Flt1), Pericyte (1,816, Rgs5 and Pdgfrb) and vascular and leptomeningeal cells VLMC (3,708, Dcn) cell types. We were unable to annotate a small group of cells and labelled them as ambiguous cell type (116) **(Supplementary Fig. 10c)**.

To further investigate tubulin gene expression changes within specific cortical cell populations, we performed pseudobulk differential expression analysis of the snRNA-seq dataset using DESeq2. Interestingly, in contrast to the predominantly decreased tubulin transcript levels observed in bulk RNA-seq, several tubulin genes exhibited increased nuclear transcript abundance in specific cell populations. Tubb3, which was significantly downregulated in bulk cortical tissue, showed increased nuclear expression in layer 2/3, layer 4, and layer 5/6 excitatory neuronal populations. Tubb4a exhibited increased nuclear expression in several neuronal populations, including immature excitatory neurons, inhibitory neurons, and postmitotic neurons, as well as oligodendrocyte precursor cells and oligodendrocytes (**Supplementary Fig. 10d; Supplementary Table 8**). These findings reveal cell-type-specific differences in nuclear tubulin transcripts that contrast with the bulk RNA-seq results, suggesting more complexity in the relationship between TTC5-mediated autoregulatory feedback and tubulin homeostasis in the developing brain.

The Gene Set Enrichment Analysis (GSEA) with hallmark datasets shows widespread changes across neuronal and non-neuronal cell types **(Supplementary Fig. 10e)**. Notably, the MYC Targets V1 gene set exhibited negative enrichment in immature excitatory neurons, neuronal progenitors, and cycling Dlx+ inhibitory progenitors. Similarly, E2F Targets and G2M Checkpoint gene sets are downregulated in astrocytes and microglia. DNA repair genes also showed negative enrichment in neuronal progenitors. In contrast, several signaling and cell-cycle-associated pathways exhibited positive enrichment in other cortical cell populations, including the Mitotic Spindle gene set in postmitotic neurons. Together, these findings suggest widespread but cell-type-dependent alterations in transcriptional programs associated with cell-cycle regulation and differentiation in the *TTC5^R263*/R263*^* cortex.

### Reduced tubulin proteins and polymerized β3-tubulin in *TTC5^R263*/R263*^* brains

In our transcriptome data, we observed downregulation of tubulin mRNAs in bulk RNA-sequencing from neonatal cortex. To assess the impact of the *TTC5^R263*/R263*^* mutation on tubulin protein levels in brain in vivo, we performed western blot analysis on mutant mouse brains, to quantify α-tubulin, β3-tubulin, acetyl-α-tubulin, γ-tubulin and pan-β-tubulin. We first looked at the α- and β-tubulin protein levels as they both form the basic building blocks of microtubules. β3-tubulin is an abundant and neuron-specific isotype of β-tubulin, which influences microtubule dynamics and stability.^29,31,86,87^ In the *TTC5^R263*/R263*^*brain, there was an approximate 30.5% reduction in α-tubulin protein levels (**Fig. 6a, b**) and a ∼38.5% reduction in β3-tubulin (**Fig. 6c, d**). We also evaluated post-translationally modified forms of α-tubulin. While α-tubulin Lys40 acetylation is not directly involved in polymerization, acetylation marks long-lived, stable microtubules, enhances microtubule (MT) flexibility and prevents microtubules (MTs) from aging characterized by structural damage.^88–90^ An approximately 30.3% decrease in acetylated α-tubulin was observed in *TTC5^R263*/R263*^*cortex. No significant differences were observed in the acetylated α-tubulin/total α-tubulin ratio, indicating that the observed reduction in acetylated α-tubulin may reflect decreased total α-tubulin rather than detectable changes in relative α-tubulin acetylation (**Supplementary Fig. 11**). Though γ-tubulin does not polymerize into microtubules directly but is essential for nucleating and assembling^91,92^, an approximate 30.3% reduction in γ-tubulin was observed in *TTC5^R263*/R263*^* cortex (**Fig. 6e, f**). Not all β-tubulin isoforms showed reduced expression at the mRNA level. Consistent with this, at the protein level, only a modest reduction of approximately 16.5% was detected in pan-β-tubulin levels (**Fig. 6g, h**). No significant differences between WT and HET genotypes were observed. Overall, these results revealed a strong reduction of tubulin protein levels in the *TTC5^R263*/R263*^*brain in vivo. Interestingly, when we observed a steady reduction of TTC5 protein across postnatal stages from P0 to 20 weeks from total brain protein in wild-type animals, a reduction of β3-tubulin was also observed (**Supplementary Fig. 12**).

Microtubules are polymers composed primarily of α-tubulin and β-tubulin heterodimers. Microtubule formation depends on tubulin availability, and reduced levels of free tubulin can limit microtubule assembly and stability.^93,94^ Therefore, we examined the polymerization state of microtubules by performing a microtubule polymerization and co-sedimentation assay in *TTC5^R263*/R263*^*and control mouse brains at P0. We observed significantly reduced levels of β3-tubulin in the polymerized fraction (P2) in *TTC5^R263*/R263*^* mutants, whereas the soluble fraction (S2) showed no significant difference (**Supplementary Fig. 13a, b**). To account for reduced total β3-tubulin abundance in *TTC5^R263*/R263*^* brains, we further normalized the P2 fraction of β3-tubulin to the corresponding input fraction (S1). The P2/S1 ratio was reduced by about 50% (mutant 33.33 ± 23.69% vs. control 64.93 ± 27.85%), although the difference was not statistically significant (**Supplementary Fig. 13c**). Together, these findings demonstrate reduced tubulin protein levels and decreased polymerized β3-tubulin in TTC5-deficient brains, consistent with disrupted tubulin homeostasis during cortical development.

### *TTC5^R263*/R263*^* mutation reduces microtubule growth dynamics and neuronal arborization in cortical neurons in vitro

To investigate the effect of the *TTC5^R263*/R263*^* mutation on growing neuronal MTs, we studied microtubule growth behavior in primary neurons from littermate control and *TTC5^R263*/R263*^*animals in vitro. We transfected primary cortical neurons at days in vitro 2 (DIV2) with EB3-tdTomato and examined live neurons with confocal microscopy at DIV3. EB3-tdTomato specifically labels the plus ends of polymerizing MTs. For detection of individual EB3-tdTomato growth tracks, we included only comets that could be followed clearly for three time frames or more (**Fig. 7a-i**). We analyzed and measured the three major parameters of MT dynamics, including comet growth rate, comet lifetime and comet growth length using ImageJ Plugin MTrackJ for motion tracking and analysis. EB3-tdTomato comet velocity was decreased by 21% (mutant 0.0873 ± 0.0184 *μ*m/sec vs. control 0.1105 ± 0.0185 *μ*m/sec), indicating that the *TTC5^R263*/R263*^* mutation impairs microtubule growth rate (**Fig. 7j**). EB3-tdTomato comet growth length, reflecting the distance of MT growth prior to pause or catastrophe, was reduced by 34% (mutant 2.594 ± 0.829 *μ*m vs. control 3.926 ± 0.761 *μ*m) (**Fig. 7l**). In contrast, EB3-tdTomato comet lifetime was not changed (mutant 6.30 ± 0.23 sec vs. control 6.40 ± 0.16 sec) (**Fig. 7k**). Together, these results suggest that the *TTC5^R263*/R263*^* mutation impairs neuronal MT growth dynamics.

We then investigated the impact of the *TTC5^R263*/R263*^*mutation on neuronal arborization. We performed continuous live-cell imaging using the IncuCyte S3 Live-Cell Analysis System (Essen BioScience) to measure the neuronal branch points and neurite length. The phase-contrast images were collected every 6 hours continuously for approximately 14∼15 DIV in total using a 10× objective lens. Image analysis was performed using the NeuroTrack image acquisition module of the IncuCyte software, which automatically identifies cell bodies and neurite processes to quantify total neurite length and the number of branch points per cell body cluster. All values were normalized to baseline measurements at time 0 h, and data are presented as fold change relative to this baseline (set to 1). As shown in **Fig. 7m, n**, *TTC5^R263*/R263*^* neurons exhibited significant reduction in both neurite length and branching points, suggesting that the *TTC5^R263*/R263*^* mutation affects neurite branching. No significant differences were observed between WT and HET genotypes. Using this automated imaging platform, real-time images were generated throughout the culturing of neurons for 14∼15 DIV (**Supplementary Fig. 14**). Therefore, in summary, the *TTC5^R263*/R263*^* mutation appears to reduce the dynamic growth behavior of microtubules in primary neurons and lead to defects in neuronal arborization.

## Discussion

Biallelic mutations in TTC5 are associated with the newly discovered neurodevelopmental syndrome, NEDCAFD, which shares clinical and neuroanatomical features with human tubulinopathies characterized by cortical malformation.^9^,^11^ Here, using a mouse model harboring the patient-derived pathogenic nonsense mutation c.787C>T; p.(Arg263Ter), we demonstrate that TTC5 is essential for normal cortical development, providing strong construct validity as a model of NEDCAFD. Germline loss of TTC5 resulted in neonatal lethality, reduced brain size, abnormal cortical development, and depletion of neural progenitor populations. These abnormalities were accompanied by altered mitotic progression, increased cell-cycle exit, changes in progenitor fate, and increased DNA damage signaling. TTC5 has been characterized as a key component of tubulin autoregulatory machinery, a process exquisitely controls tubulin availability to support microtubule function.^23–27^ We further identified widespread alterations in tubulin transcript and protein abundance, along with impaired microtubule growth dynamics and reduced neuronal arborization.

The TTC5 nonsense variant c.787C>T, p.(Arg263Ter), is the most frequently reported pathogenic variant among affected individuals with NEDCAFD.^9,11^ This mutation is located between the last tetratricopeptide repeat (TPR) domain and the oligonucleotide binding (OB) domain.^9,11^ Our data support Arg263Ter as a strong loss-of-function allele, with substantially reduced full-length TTC5 transcript and no detectable full-length or truncated TTC5 protein in homozygous mutant brains.

The expansion of the cerebral cortex depends on coordinated proliferation and differentiation of NPCs. Although the common and multiple progenitor models propose distinct mechanisms of cortical neuronal lineage specification, both emphasize the importance of robust progenitor amplification in establishing cortical size and cellular diversity.^59, 95–97^ We show that TTC5 is required for normal cortical progenitor expansion during development. In *TTC5^R263*/R263*^* embryos, we observed pronounced reductions in both radial glia and intermediate progenitors during mid-neurogenesis, accompanied by persistent reductions in proliferating neural progenitors at birth. These abnormalities were associated with reduced cortical thickness and a more significant reduction in superficial cortical neuronal populations. Given the sequential generation of cortical layers, these findings are consistent with the greater dependence of later-born neurons on sustained progenitor expansion throughout corticogenesis.

The reduction in cortical progenitor populations was accompanied by altered mitotic behavior and cell fate dynamics. At E14.5, *TTC5^R263*/R263*^* RGCs exhibit increased mitotic index and accumulation of cells in prometaphase and metaphase, consistent with delayed progression through these mitotic stages. We observed altered cleavage-plane orientation, with fewer vertical divisions and increased horizontal and oblique divisions, which could contribute to abnormal progenitor behavior. EdU labeling at E13.5 provided additional evidence of altered progenitor dynamics, revealing increased cell-cycle exit and a greater proportion of EdU-labeled cells expressing the early neuronal marker Tbr1, accompanied by reduced proportions expressing progenitor markers Pax6 and Tbr2. Together, these findings support a shift toward neuronal differentiation at the expense of progenitor maintenance, which could progressively limit progenitor expansion and contribute to reduced cortical growth.

Our transcriptomic analyses further identified altered expression of cell-cycle-associated genes that may contribute to progenitor abnormalities. Bulk RNA-seq revealed decreased expression of G1/S regulators, including CDK4, CDK2, and associated cyclins, alongside increased expression of cyclin-dependent kinase inhibitors. Although tissue-level changes may partly reflect altered cortical cellular composition, pseudobulk snRNA-seq analysis identified similar expression differences within defined progenitor populations, including reduced CDK4 and CDK6 and increased CDKN1A expression. Cell-type-specific gene set enrichment analysis further revealed alterations in proliferation-associated pathways, including reduced enrichment of MYC targets in neuronal progenitors. These data suggest that changes in cell-cycle regulatory pathways may contribute to impaired progenitor expansion. Whether TTC5 deficiency directly disrupts cell cycle progression or other interphase regulatory mechanisms remains to be determined.

TTC5 deficiency is associated with profound alterations in tubulin abundance and microtubule function in the developing cortex. TTC5 has been characterized as a key component of tubulin autoregulation, a negative-feedback mechanism regulating tubulin mRNA in response to soluble tubulin availability.^23–25^ Disruption of this mechanism might be expected to increase tubulin mRNA abundance. Interestingly, our bulk RNA-seq analysis revealed decreased expression of multiple tubulin transcripts in neonatal *TTC5^R263*/R263*^* cortex, accompanied by reductions in several tubulin proteins, including α-tubulin, β3-tubulin, γ-tubulin, acetylated α-tubulin, and, to a lesser extent, pan-β-tubulin. Altered cellular composition in the mutant cortex could contribute to tissue-level changes in tubulin transcript and protein levels. This is particularly relevant for β3-tubulin, a neuron-enriched tubulin isotype, given the altered neuronal populations observed in *TTC5^R263*/R263*^* cortices. The consistent reductions across multiple tubulins suggest widespread disruption of tubulin abundance in the TTC5-deficient brain, although the extent to which these changes reflect cell-intrinsic alterations remains an important question. Cell intrinsic mechanisms appear to be supported by other studies in neurons in vitro, where TTC5 loss-of-function has been shown to decrease tubulin transcript and protein levels in neuronal cell lines and iPSC-derived neuronal culture, including β3-tubulin.^10,98^

Our single-nucleus transcriptomic analyses further revealed distinct patterns of tubulin expression across cortical cell populations. In contrast to the predominantly decreased tubulin transcript abundance observed in bulk RNA-seq, pseudobulk analysis identified increased nuclear abundance of several tubulin transcripts, including Tubb3 in cortical excitatory neurons and Tubb4a in multiple neuronal and oligodendrocyte-lineage populations. Given that TTC5-dependent autoregulation operates on translating tubulin mRNAs in the cytoplasm, one possibility is that increased nuclear abundance of selected tubulin transcripts represents a compensatory response to reduced tubulin availability. Alternatively, the discrepancy may arise from differences in cellular composition, RNA processing, nuclear export, or transcript stability. These findings suggest a more complex relationship between TTC5-dependent autoregulatory feedback and tubulin nuclear gene expression in the developing brain, potentially involving additional levels of regulation beyond cytoplasmic mRNA degradation alone.

In addition to altered tubulin abundance, TTC5 deficiency was associated with impaired microtubule function. Microtubule co-sedimentation assays showed reduced polymerized β3-tubulin in mutant brain lysates. Importantly, live-cell imaging of EB3-labeled microtubules in primary cortical neurons further demonstrated decreased microtubule growth velocity and growth length without significant changes in comet lifetime, indicating impaired microtubule growth dynamics. These abnormalities were accompanied by reduced neurite length and branching, consistent with the critical role of microtubule dynamics in neuronal morphogenesis.

The convergence of progenitor mitotic abnormalities and altered microtubule biology points to a potential mechanistic relationship between these phenotypes. Disrupted tubulin availability and microtubule dynamics could compromise mitotic spindle function, contributing to abnormal mitotic progression and altered progenitor behavior.^69–72^ Furthermore, prolonged mitosis, particularly delayed prometaphase progression, has been shown to bias radial glial progenitors toward premature neuronal differentiation or cell-cycle exit.^84,85^ Altered tubulin regulation and microtubule function may contribute to the mitotic and progenitor abnormalities observed in *TTC5^R263*/R263*^* cortex. Nevertheless, a direct causal relationship remains to be established.

TTC5 has also been implicated in DNA damage response (DDR) signaling through its regulation of p53 activity.^21^ We observed increased γ-H2AX foci in *TTC5^R263*/R263*^* cortical progenitors, together with altered enrichment of DNA repair-associated pathways in neuronal progenitors. These changes may reflect consequences of abnormal mitotic progression, as mitotic perturbations have been associated with increased DNA damage.^78–80^ Alternatively, disrupted TTC5-dependent DDR signaling could independently contribute to the observed progenitor abnormalities. Notably, increased DNA damage signaling was not accompanied by detectable increases in apoptosis at the developmental stages examined. Given the role of TTC5 in augmenting p53 activity, TTC5 deficiency may impair p53-dependent apoptotic responses to mitotic stress and DNA damage, while potentially favoring p53-independent cell-cycle exit or neuronal differentiation.^85^ Future studies examining p53 pathway functions in TTC5-deficient progenitors will be important for distinguishing these mechanisms.

In summary, our study establishes TTC5 as an essential regulator of cortical neurodevelopment in vivo, required for normal progenitor expansion and tubulin homeostasis in the developing brain, providing mechanistic insights into the pathogenesis of NEDCAFD.

## Acknowledgements

The authors thank Dr. Jinping Luo and the Brown University Transgenic and Genome Editing Facility for designing and generating the TTC5^R263*/R263*^ mouse model.

## Funding

This work was supported by National Institutes of Health grants R01NS121618 (E.M.M.), R01AG087455 (E.M.M.), R01NS113141 (E.M.M.).

## Author information

Z.L., L.M., and E.M.M. designed the study. Z.L. and L.M. performed experiments and data analysis. H.R. performed the snRNAseq experiment and data analysis. Q.W. analyzed the bulk RNA sequencing dataset. M.S. and M.F. performed research and acquired data. Z.L., L.M., and E.M.M. wrote and edited the manuscript. E.M.M. supervised the entire study.

## Competing interests

The authors report no competing interests.

## Data and materials availability

All data are available in the main text or the supplementary materials. Further information and requests for resources and reagents should be directed to and will be fulfilled by Eric Morrow

---

**Supplementary Fig. 1.**
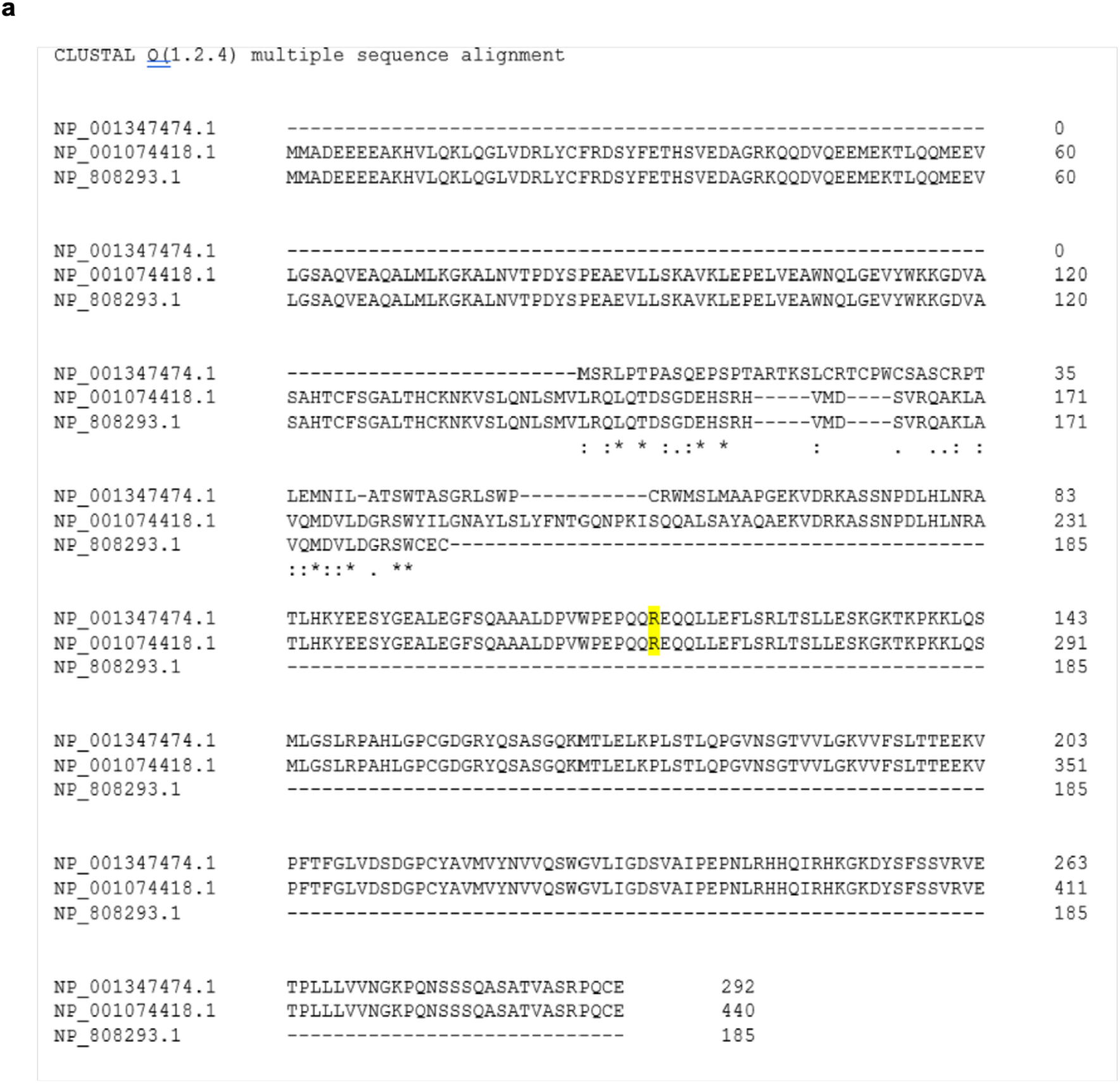
TTC5 protein CLUSTAL alignment. **a** CLUSTAL alignment of the protein sequences from all three Ttc5 transcripts. The highlighted R denotes amino acid residue R263.

**Supplementary Fig. 2.**
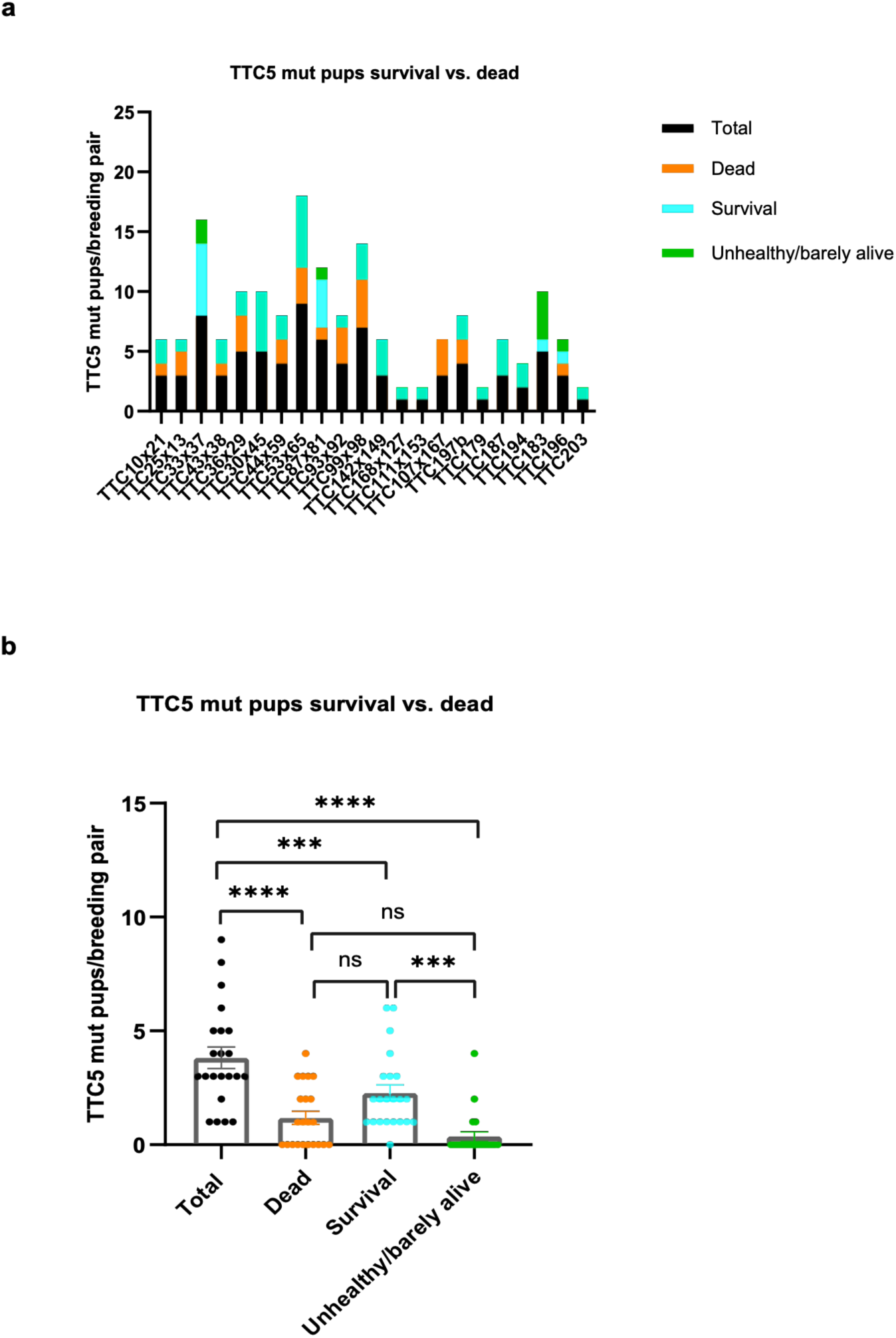
*TTC5^R263*/R263*^* leads to newborn lethality. **a** Graph exported from Graph prism shows the total mutant pups from 22 breeding pairs. Black denotes total mutant pups; orange, cyan, green indicate dead pups, survival pups, unhealthy/barely alive pups respectively newborn <p1. **b** Graph exported from Graph prism shows statistical comparison between total mutant pups, dead, survival, unhealthy/barely alive pups newborn <p1 (****, *p* < 0.0001 for total vs. dead; ***, *p =* 0.0002 for total vs. survival; ****, *p* < 0.0001 for total vs. unhealthy/barely alive; ns, *p* = 0.1140 for dead vs. survival; ns, *p* = 0.1911 for dead vs. unhealthy; ***, *p* = 0.0005 for survival vs. unhealthy/barely alive.

**Supplementary Fig. 3.**
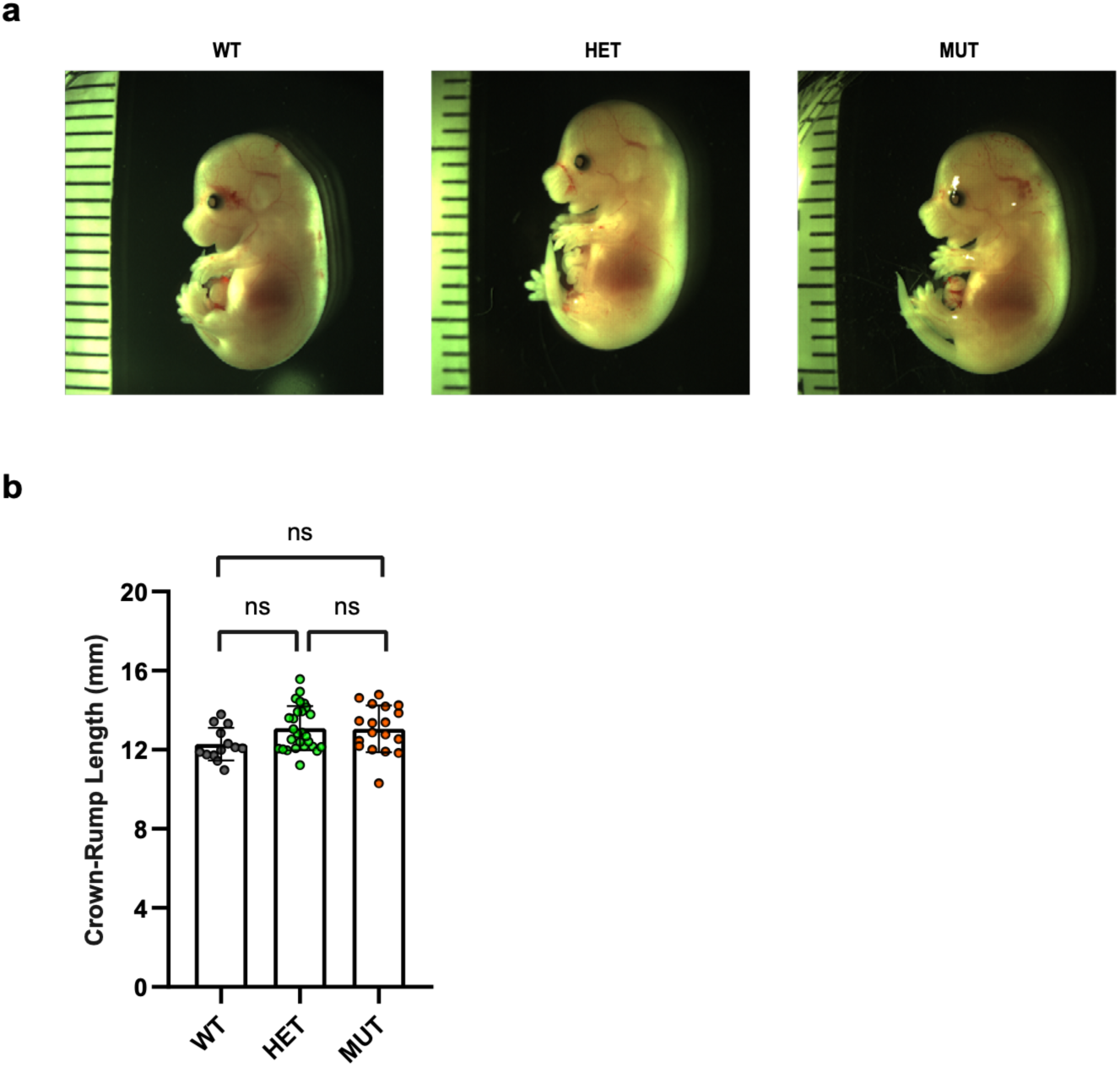
E14.5 embryos crown-rump length measurement. **a** Representative image of E14.5 embryos from the WT, HET and MUT genotypes (shown from left to right). **b** Graph exported from Graph prism showing statistical comparisons of WT, HET and MUT genotypes based on the measurement of Crown-Rump Length. One-way Anova with Tukey’s multiple comparisons test was used (ns, *p* =0.0785 for WT vs. HET; ns, *p* =0.1254 for WT vs. MUT; ns, *p* =0.9968 for HET vs. MUT). WT=13, HET=27 and MUT=18 embryos.

**Supplementary Fig. 4.**
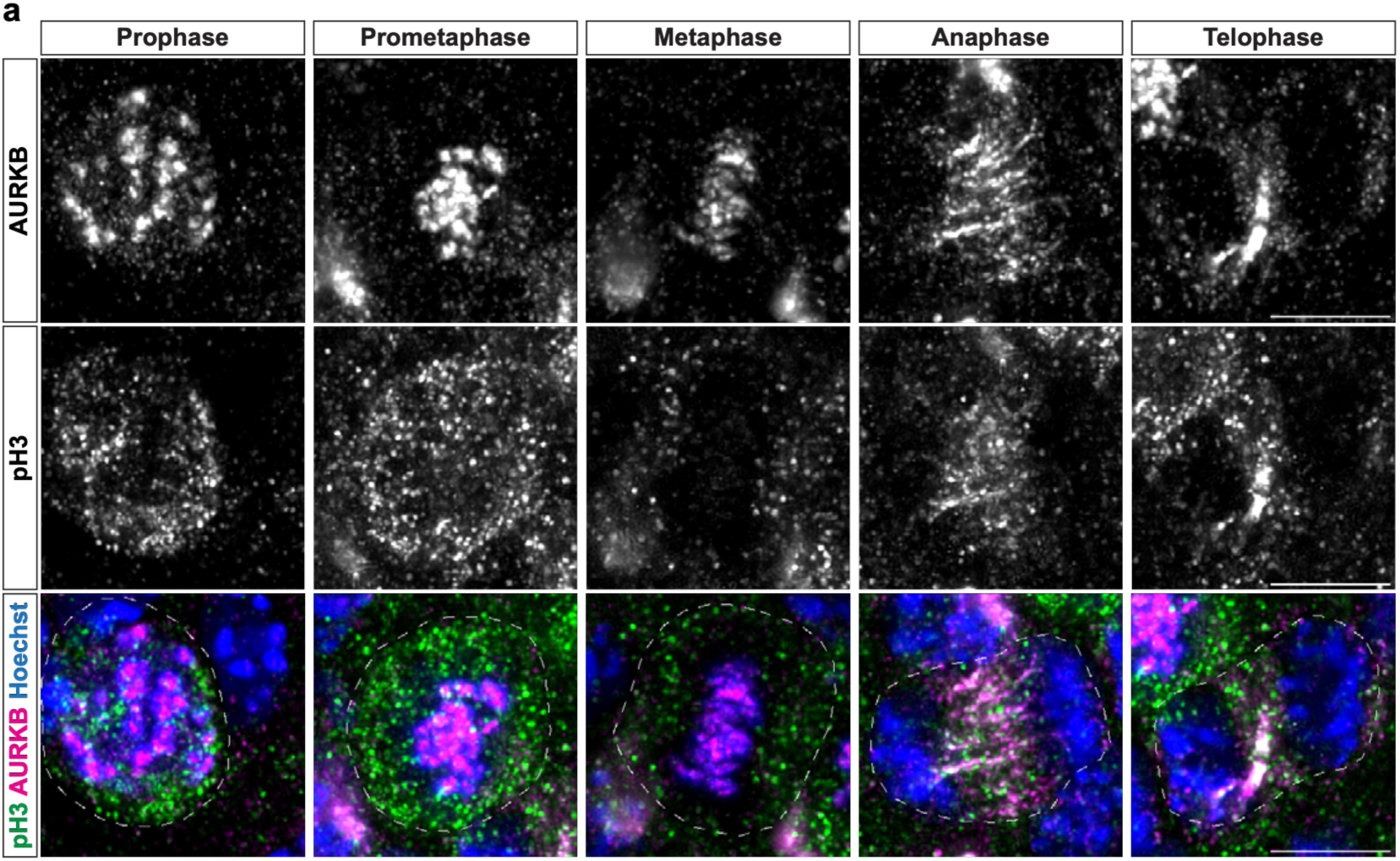
Identification of RGC mitotic stages. **a** Localization of mitotic marker phospho-Histone H3 (pH3, green) in relation to mitotic spindle marker aurora kinase B (AURKB, magenta) and Hoechst 33342 (blue) in RGCs during M phase for identification of mitotic stages (representative images from wild-type E14.5 cortical sections). During early stages of mitosis, from prophase to metaphase, AURKB is colocalized with the condensing chromosomes and pH3 is highly expressed in the nucleus. In late stages, anaphase to telophase, AURKB and pH3 are colocalized at the central spindle and the midbody region. Scale bars, 5m.

**Supplementary Fig. 5.**
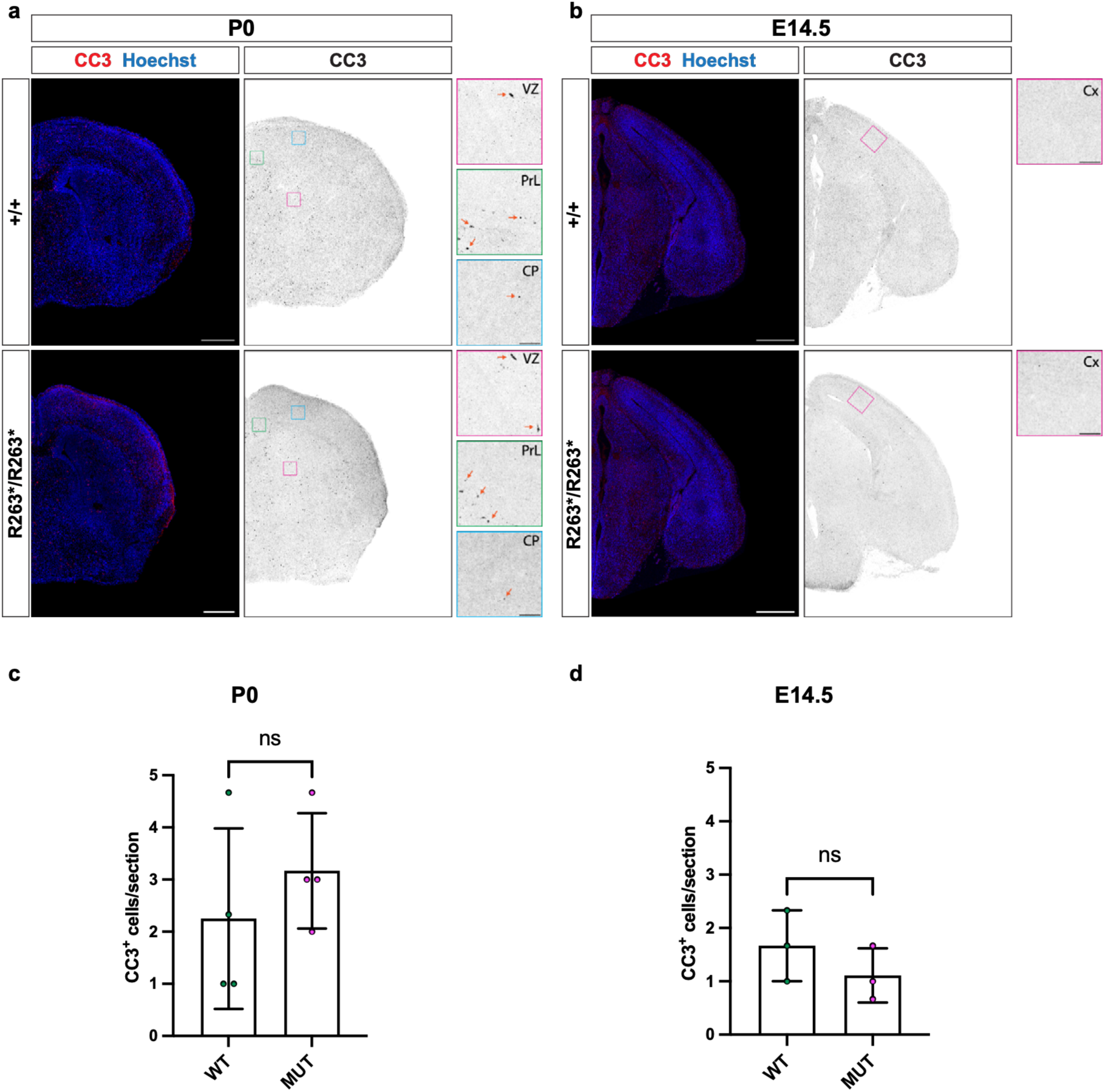
Apoptosis detection by cleaved caspase-3 immunostaining in the developing cortex. **a** P0 and **b** E14.5 wild type and TTC5*^R263*/R263*^* brain sections labeled with cleaved caspase 3 (CC3, red), a marker for apoptosis. Rectangle areas are shown in higher magnification on the right in respective colors. Orange arrows depict CC3 positive cells. VZ, ventricular zone; PrL, prelimbic cortex; CP, cortical plate; Cx, cerebral cortex. **c, d** Quantification of CC3^+^ cells in the P0 neuroepithelium (ns, *p* = 0.41) and E14.5 developing cortex (ns, *p* = 0.32) showed no significant change in the number apoptotic cells. Scale bars, 500μm, 5 μm. Sections were counterstained with Hoechst 33342 (blue) to label nuclei. n_P0_ = 4 WT and 4 MUT animals. n_E14.5_ = 3 WT and 3 MUT animals. Data are presented as means ± SDs. Statistical analyses were conducted using two-tailed t test with Welch’s correction.

**Supplementary Fig. 6.**
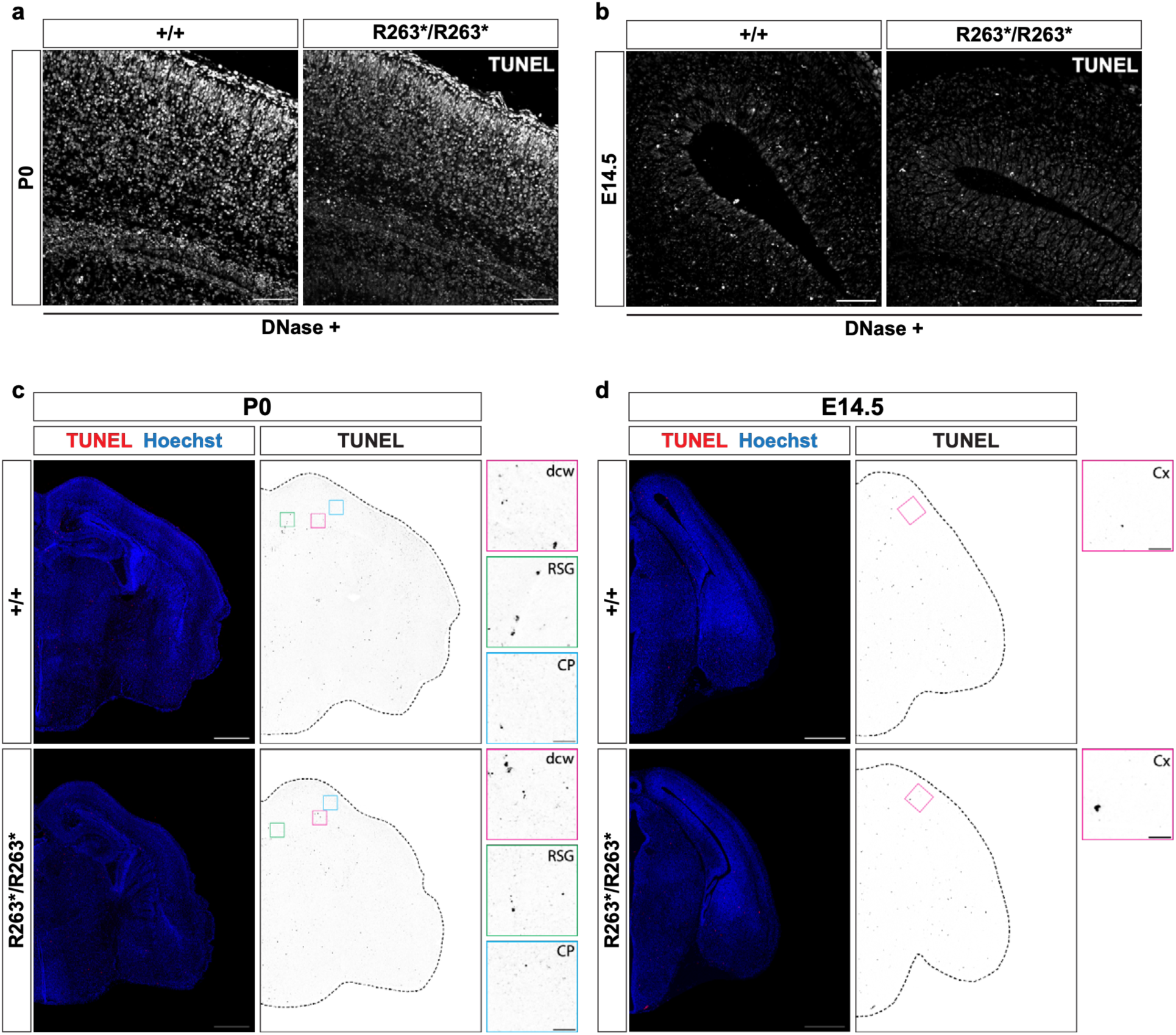
Cell death detection by TUNEL assay in the developing cortex. P0 and E14.5 wild type and TTC5*^R263*/R263*^*brain sections labeled with TUNEL assay for DNA fragmentation in cell death, with (**a, b**) or without (**c, d**) DNase treatment. dcw, deep cerebral white; RSG, retrosplenial granular cortex. Scale bars, 500μm, 50μm. Sections were counterstained with Hoechst 33342 (blue) to label nuclei. n_P0_ = 4 WT and 4 MUT animals. n_E14.5_ = 3 WT and 3 MUT animals.

**Supplementary Fig. 7.**
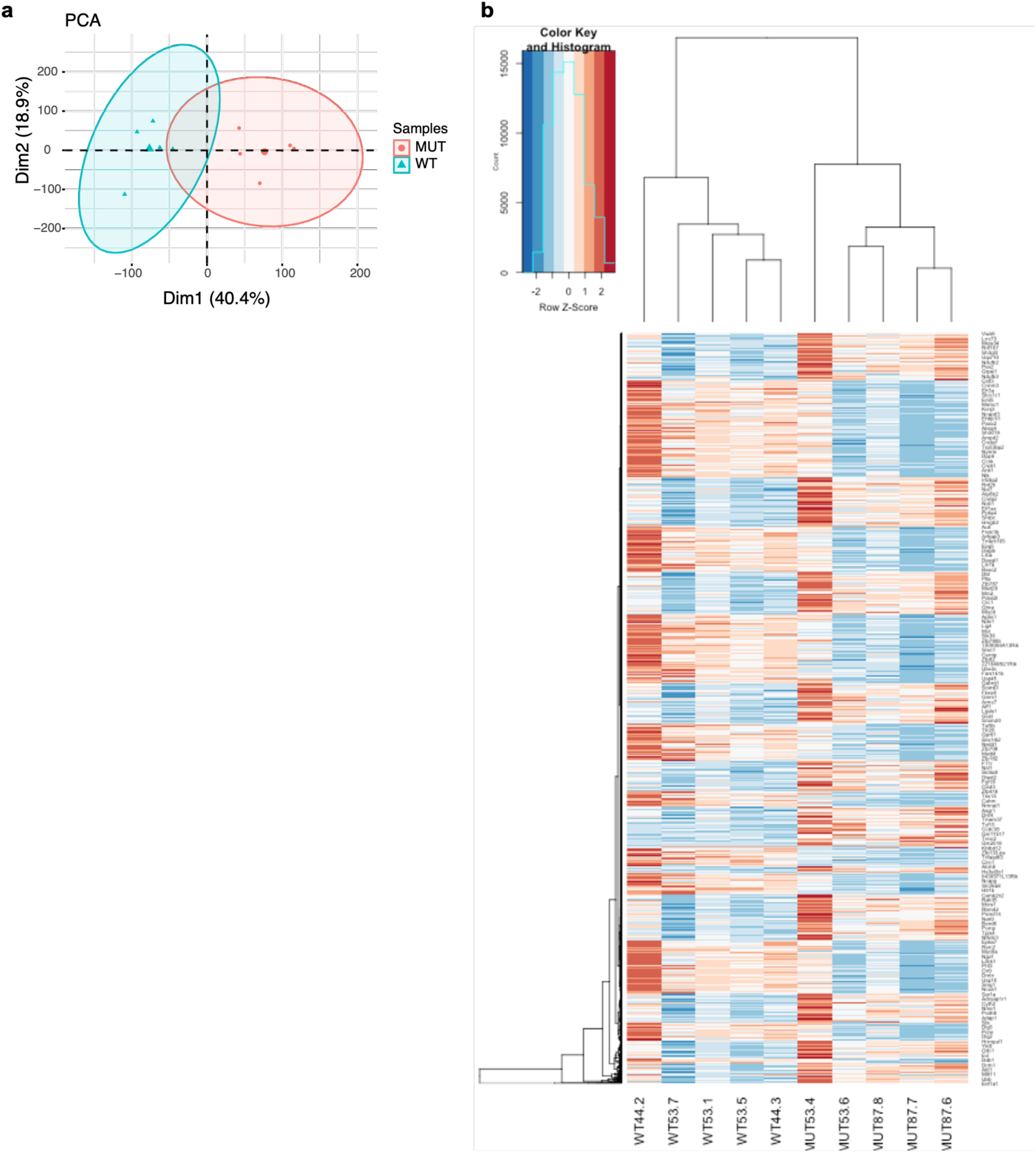
TTC5*^R263*/R263*^* bulk RNA sequencing PCA plot and DEGs. **a** PCA plot shows the major sample variation between WT and MUT**. b** Heatmap using analyzed DEGs expression data

**Supplementary Fig. 8.**
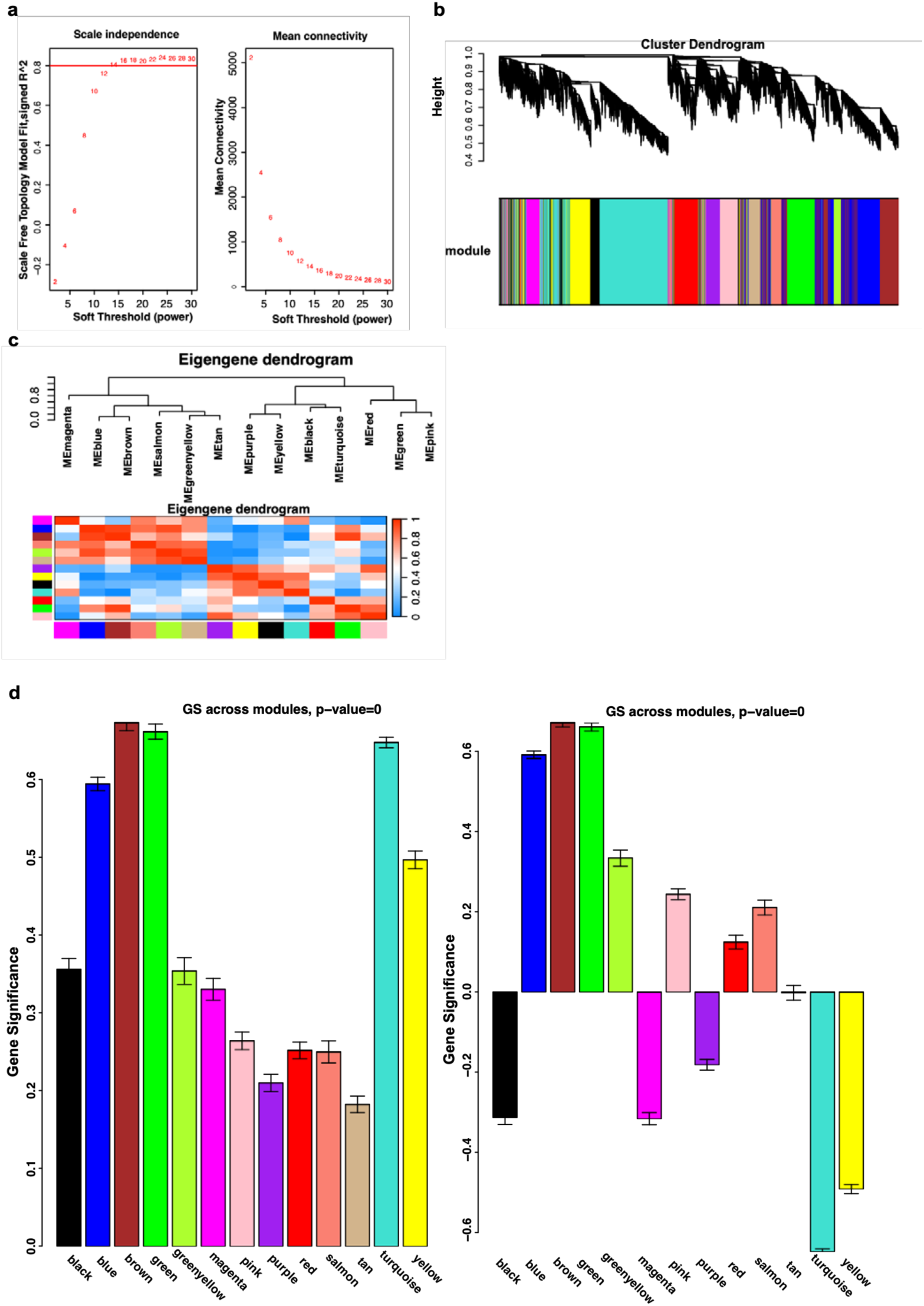
TTC5*^R263*/R263*^* bulk RNA sequencing WGCNA. **a** Scale independence (left panel) and Mean connectivity (right panel). **b** Hierarchical clustering based on gene expression distance and then using dynamic “hybrid” tree cut algorithm to cut the dendrogram with a setting of minimum cluster size at 100, and no deep split. Hierarchical clustering after merging high similar modules. **c** Module eigengene dendrogram, indicating the relationship across modules. **d** Correlation coefficient of each module to expression of sample status by barplot.

**Supplementary Fig. 9.**
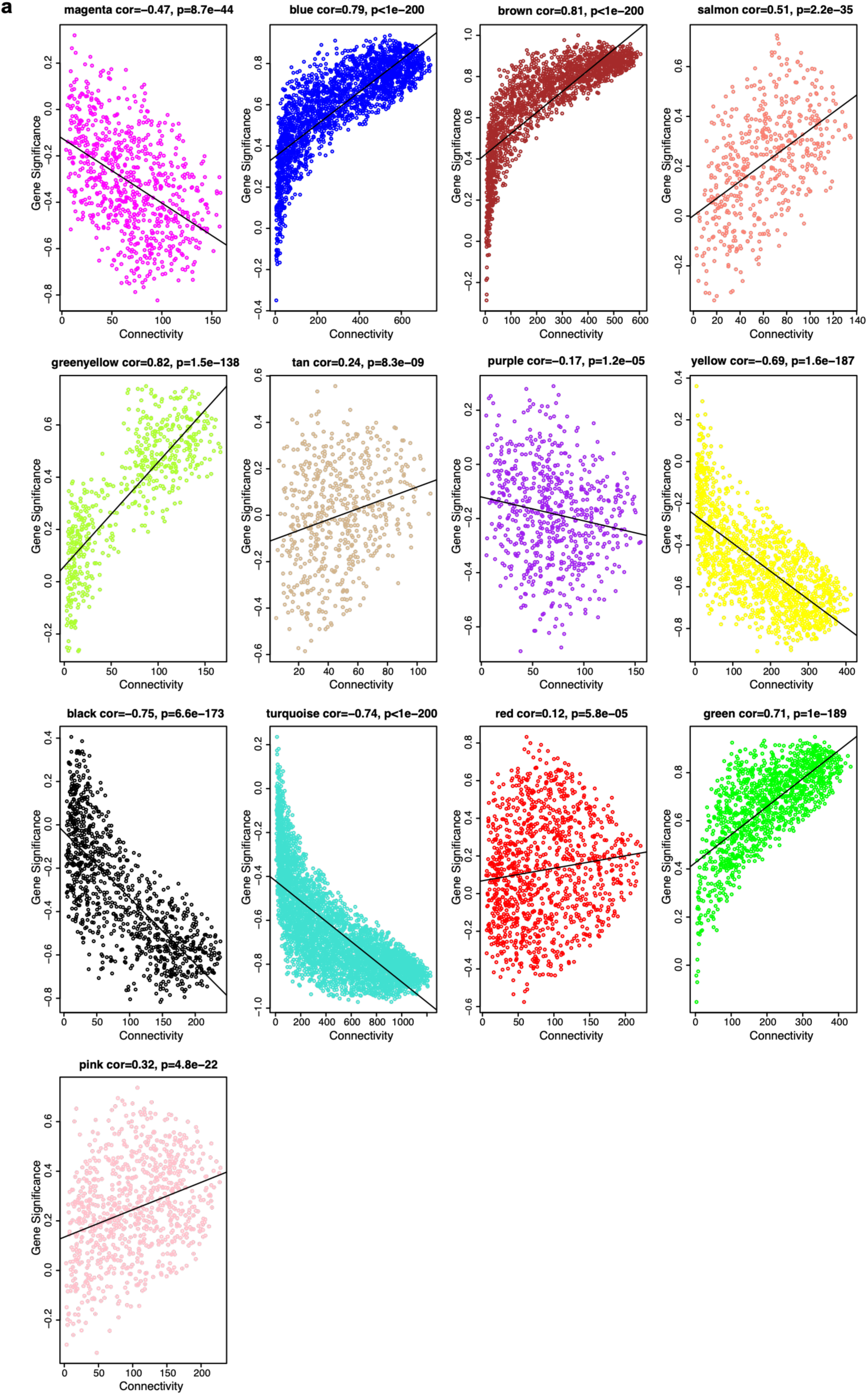
TTC5*^R263*/R263*^*bulk RNA sequencing WGCNA module correlation. **a** Correlation between module gene significance and module gene connectivity by scatterplot.

**Supplementary Fig. 10.**
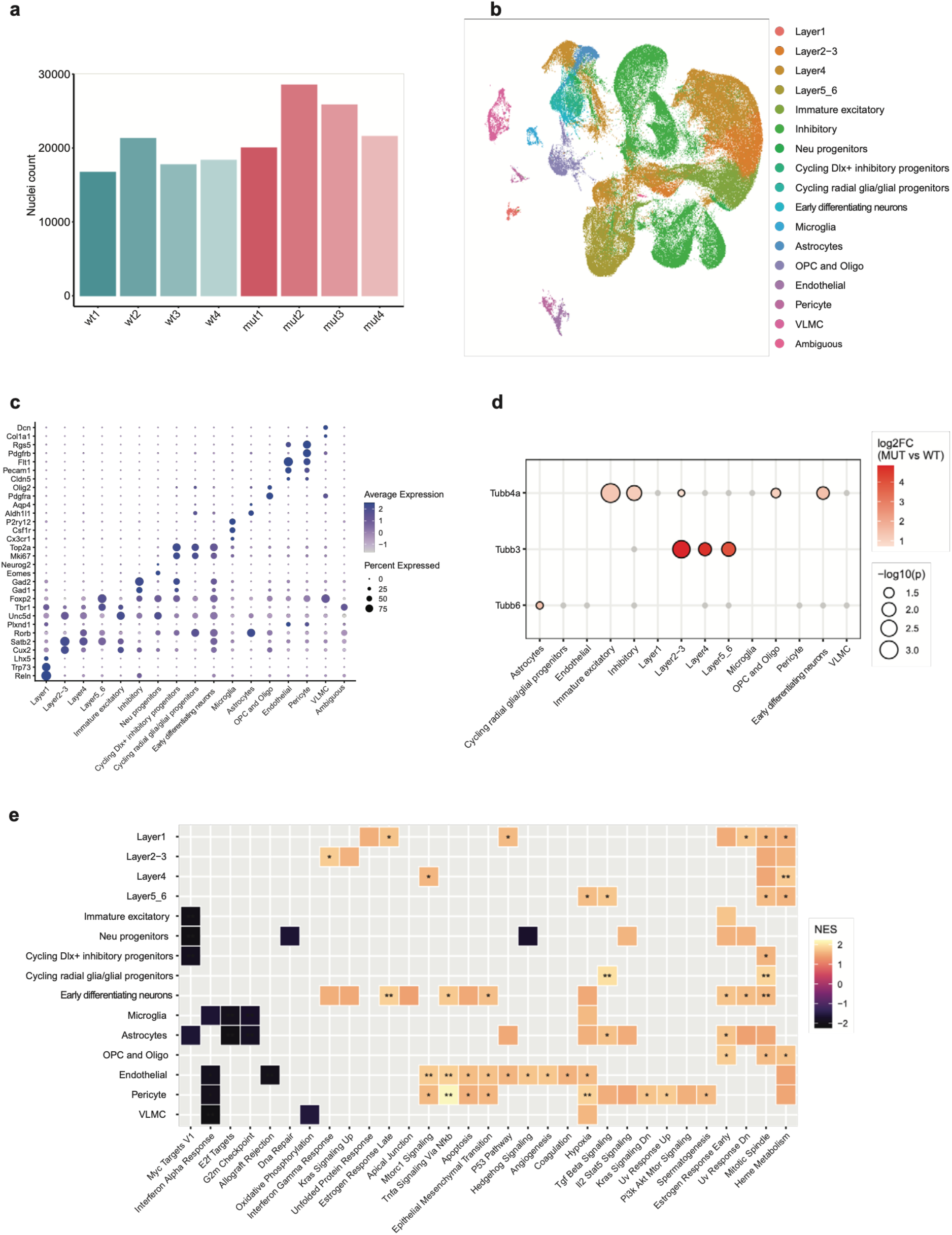
snRNA sequencing analysis of *TTC5* neocortex. **a** Total number of nuclei analyzed from each sample in the snRNA-seq. **b** Uniform Manifold Approximation and Projection (UMAP) plot shows all 174,444 nuclei (WT; 76,504, MUT 97,940) included for analysis. Clustering analysis identified 16 distinct major cell types. **c** Dotplot shows individual marker genes for each cell type. **d** Dotplot shows DEGs of β-tubulin genes (Tubb3, Tubb4a, Tubb6) across cell types from pseudobulk scRNAseq analysis (n = 4 WT, 4 MUT per cell type; DESeq2 Wald test). Colored dots indicate a pvlaue < 0.05. grey color shows no significance. **e** Heatmap shows gene set enrichment (GSEA) analysis results from Hallmark pathways across cell types. Colors indicate normalized enrichment scores (NES). Padj < 0.1 (color), * padj < 0.05, ** padj < 0.01, grey color shows not significant. n = 4 WT (3 males and 1 females) and 4 MUT (2 males and 2 females) animals from 3 litters.

**Supplementary Fig. 11.**
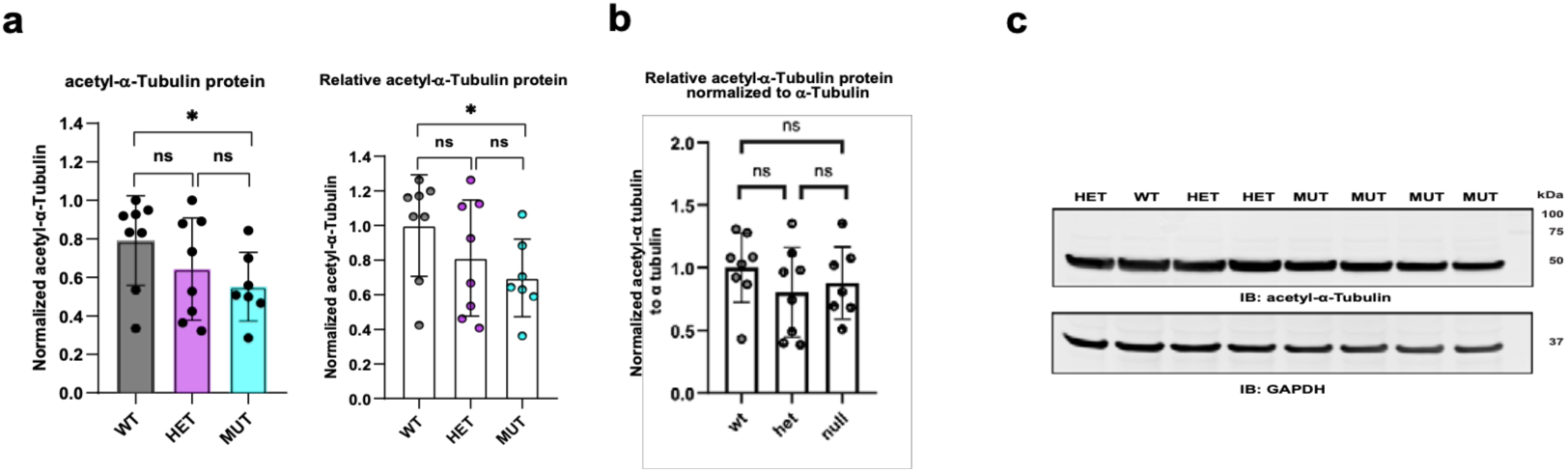
Acetyl-α -tubulin protein expression in *TTC5^R263*/R263*^* brains. **a** Quantification and comparison of acetyl-α -tubulin between *TTC5^+/+^*, *TTC5^+/R263*^*, and TTC5*^R263*/R263*^* animals (*, *p* = 0.0418 for WT vs. MUT; ns, *p* = 0.4455 for HET vs. MUT; ns, p=0.2531 for WT vs. HET). **b** Quantification and comparison of acetyl-α -tubulin normalized to α -tubulin between *TTC5^+/+^*, *TTC5^+/R263*^*, and TTC5*^R263*/R263*^*animals (ns, *p* = 0.7279 for WT vs. MUT; ns, *p* = 0.8934 for HET vs. MUT; ns, p=0.4323 for WT vs. HET). **c** Representative gel image for acetyl-α -tubulin is shown. One-way Anova with Tukey’s multiple comparisons test. N = 8 WT (3 males and 5 females), 8 HET (6 males and 2 females) and 7 MUT (2 males and 5 females) animals. Data are presented as means ± SDs.

**Supplementary Fig. 12.**
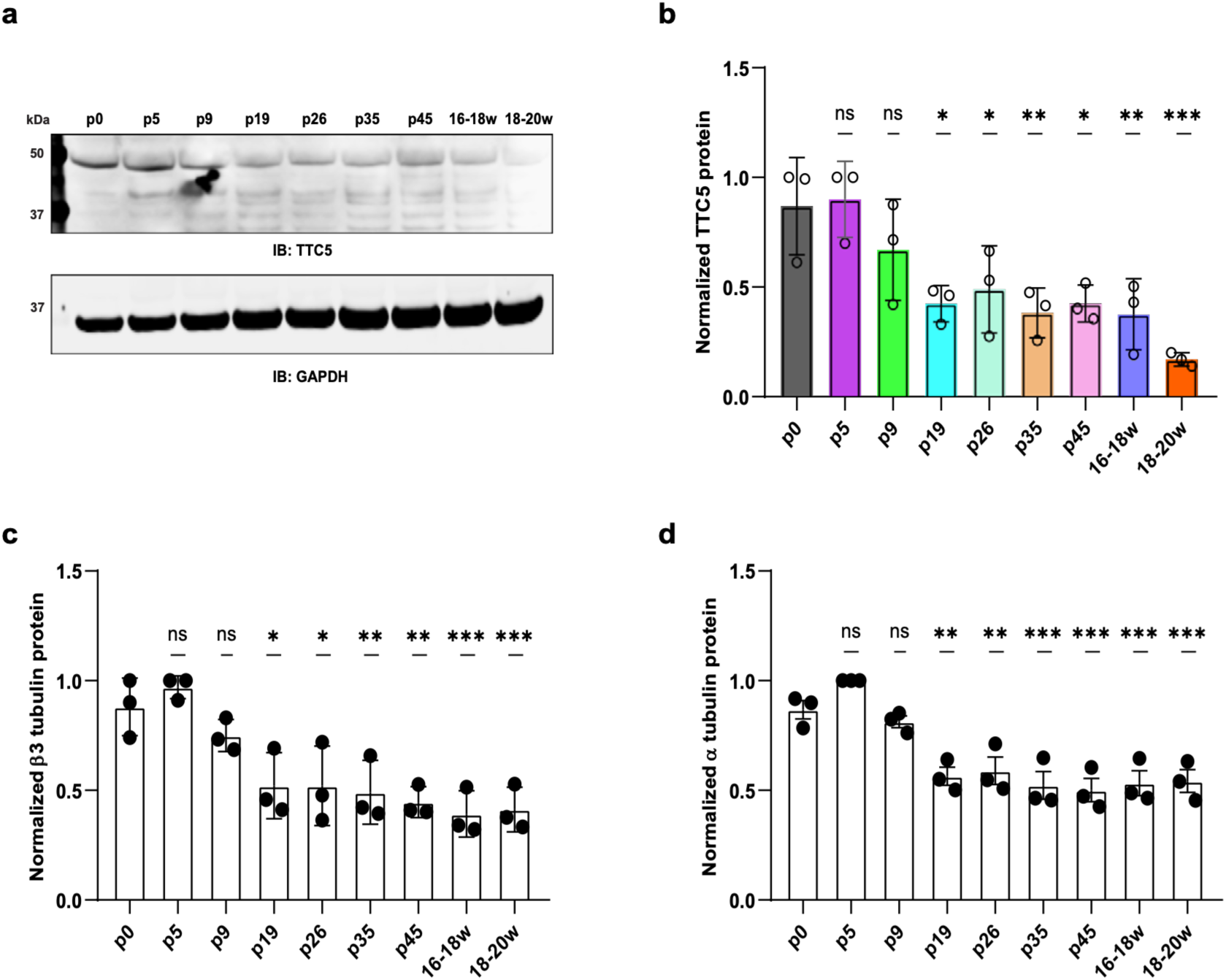
TTC5 and β3-tubulin protein expression across postnatal stages. **a** Western blotting was performed to determine TTC5 protein expression from Swiss Webster mouse brain tissue homogenates harvested from postnatal P0, -5, -9, -19, -26, -35, -45, ∼16-18w and ∼18-20w. GAPDH was used as control. **b** TTC5 protein was normalized to GAPDH and compared between P0 with other time points. One-way Anova with Dunnett’s multiple comparisons test was used (ns, *p* =0.9997 for P0 vs. P5; ns, *p* =0.5347 for P0 vs. P9; *, *p* =0.0175 for P0 vs. P19; *, *p* =0.0496 for P0 vs. P26; **, *p* 0.0087 for P0 vs. P35; *, *p* =0.0176 for P0 vs. P45; **, *p* =0.0077 for P0 vs. 16-18w and ***, *p* =0.0003 for P0 vs. 18-20w). Graph exported from Graph prism was shown. **c** β3-tubulin protein was normalized to GAPDH and compared between P0 with other time points. One-way Anova with Dunnett’s multiple comparisons test was used (ns, *p* =0.9153 for P0 vs. P5; ns, *p* =0.6687 for P0 vs. P9; *, *p* =0.0105 for P0 vs. P19; *, *p* =0.0104 for P0 vs. P26; **, *p* =0.0055 for P0 vs. P35; **, *p* =0.002 for P0 vs. P45; ***, *p* =0.0006 for P0 vs. 16-18w and ***, *p* =0.001 for P0 vs. 18-20w). Graph exported from Graph prism was shown. **d** α -tubulin protein was normalized to GAPDH and compared between P0 with other time points. One-way Anova with Dunnett’s multiple comparisons test was used (ns, *p* =0.2989 for P0 vs. P5; ns, *p* =0.9527 for P0 vs. P9; **, *p* =0.0020 for P0 vs. P19; **, *p* =0.0044 for P0 vs. P26; ***, *p* =0.0006 for P0 vs. P35; ***, *p* =0.0003 for P0 vs. P45; ***, *p* =0.0007 for P0 vs. 16-18w and ***, *p* =0.0010 for P0 vs. 18-20w). Graph exported from Graph prism was shown.

**Supplementary Fig. 13.**
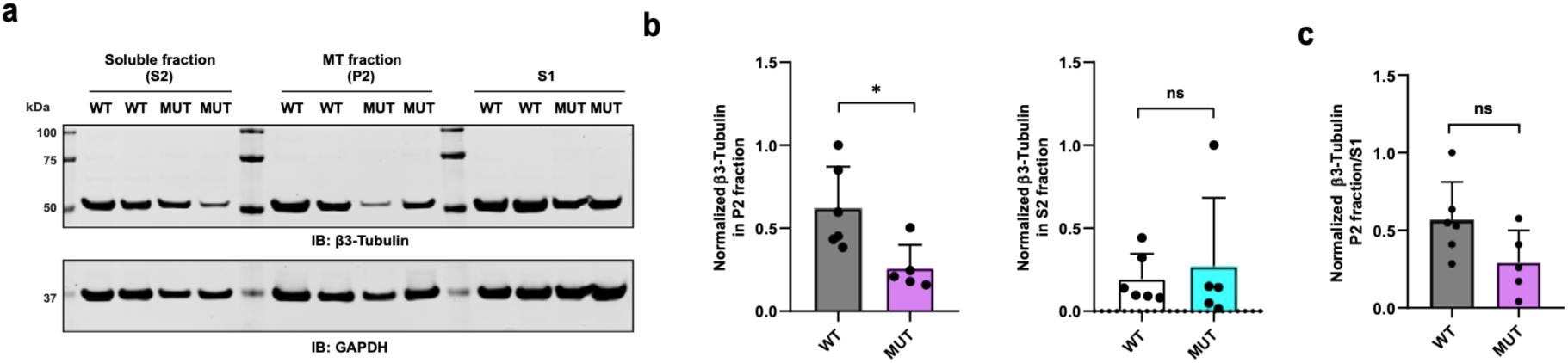
Polymerized tubulin is reduced in *TTC5^R263*/R263*^* brains. Gel images for β3-tubulin from P0 mouse brain tissue homogenized and fractionated from polymerized (P2), soluble microtubules (S2) and supernatant (S1). **b** Quantification and comparison of P2 (*, *p* = 0.0166) and S2 (ns, *p*=0.7058) fractionation. **c** Quantification and comparison of P2/S1 ratio (ns, *p*=0.0726). Unpaired two-tailed t-test with Welch’s correction. n = 6 WT (3 males and 3 females) and 5 MUT (1 male and 4 females) animals. Data are presented as means ± SDs.

**Supplementary Fig. 14.**
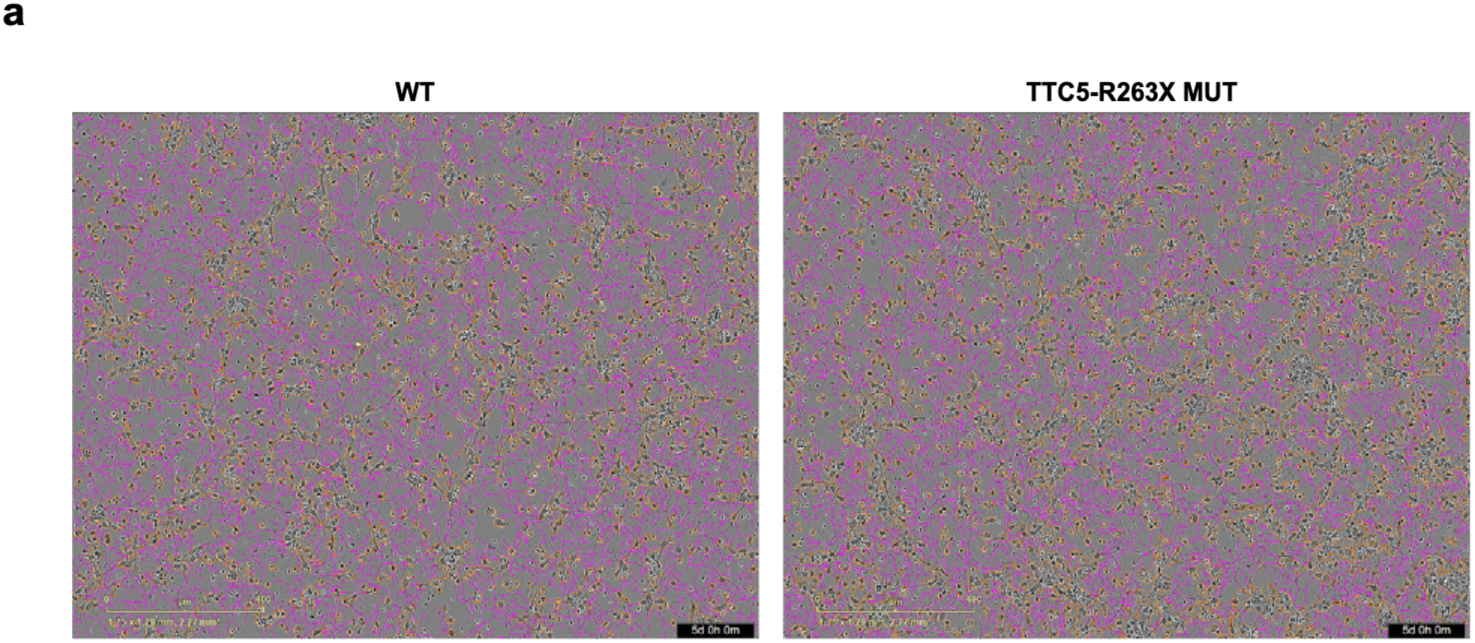
IncuCyte S3 live-cell analysis system images. **a** IncuCyte S3 live-cell analysis system image from a single well of cortical neurons showing the phase image and masking for identifying neurites (purple) and cell bodies (orange) at DIV5. Scale bar, 400 μm. Left panel: Representative image from WT; right panel: Representative image from TTC5*^R263*/R263*^*.

## Supplementary Table Legends

**Supplementary Table 1**: All genes from bulk RNA sequencing.

**Supplementary Table 2**: Differentially expressed genes in mouse Ttc5 mutant compared with wild type.

**Supplementary Table 3**: Cell cycle genes from bulk RNA sequencing.

**Supplementary Table 4:** Tubulin genes from Bulk RNA sequencing (extended from Table 1).

**Supplementary Table 5:** Enriched pathways among differentially expressed genes.

**Supplementary Table 6:** Gene list in WGCNA modules.

**Supplementary Table 7:** Enriched pathways among genes in each WGCNA modules.

**Supplementary Table 8**: All genes from snRNA sequencing pseudobulk analysis.

**Supplementary Table 9:** Primers used in this study.

## Notes

### Competing Interest Statement

The authors have declared no competing interest.

